# Neural decoding of the speech envelope: Effects of intelligibility and spectral degradation

**DOI:** 10.1101/2024.02.20.581129

**Authors:** Alexis Deighton MacIntyre, Robert P. Carlyon, Tobias Goehring

**Author notes:** Correspondence concerning this article should be addressed to Alexis Deighton MacIntyre, MRC Cognition and Brain Sciences Unit, University of Cambridge, 15 Chaucer Road, Cambridge, CB2 7EF, United Kingdom. To appear in *Trends in Hearing* (2024).

## Abstract

During continuous speech perception, endogenous neural activity becomes time-locked to acoustic stimulus features, such as the speech amplitude envelope. This speech-brain coupling can be decoded using non-invasive brain imaging techniques, including electroencephalography (EEG). Neural decoding may provide clinical use as an objective measure of stimulus encoding by the brain - for example during cochlear implant (CI) listening, wherein the speech signal is severely spectrally degraded. Yet, interplay between acoustic and linguistic factors may lead to top-down modulation of perception, thereby complicating audiological applications. To address this ambiguity, we assess neural decoding of the speech envelope under spectral degradation with EEG in acoustically hearing listeners (n = 38; 18-35 years old) using vocoded speech. We dissociate sensory encoding from higher-order processing by employing intelligible (English) and non-intelligible (Dutch) stimuli, with auditory attention sustained using a repeated-phrase detection task. Subject-specific and group decoders were trained to reconstruct the speech envelope from held-out EEG data, with decoder significance determined via random permutation testing. Whereas speech envelope reconstruction did not vary by spectral resolution, intelligible speech was associated with better decoding accuracy in general. Results were similar across subject-specific and group analyses, with less consistent effects of spectral degradation in group decoding. Permutation tests revealed possible differences in decoder statistical significance by experimental condition. In general, while robust neural decoding was observed at the individual and group level, variability within participants would most likely prevent the clinical use of such a measure to differentiate levels of spectral degradation and intelligibility on an individual basis.

## Introduction

Auditory speech perception is made possible by the continuous interaction between cognitive processes, including learning, memory, and motor knowledge acquired from speech production, and sensory encoding of the acoustic signal at hierarchically organised levels (Davis & Johnsrude, 2007). Despite this complexity, the neural signature of some aspects of speech perception can be detected using non-invasive brain imaging methods, such as electroencephalography (EEG). Whereas these tools have largely been used to examine the brain response to short, isolated speech sounds, such as syllables or words, listeners’ neural activity also covaries with continuous speech features, including the amplitude envelope or phonetic annotations, across time, thereby enabling the use of engaging, ecologically relevant stimuli (e.g., audiobook recordings; Lalor & Foxe, 2010; Di Liberto et al., 2015; Ding & Simon, 2014; Golumbic et al., 2013). This speech-brain correspondence can be recovered using both linear and nonlinear methods, such as cross-correlation (Aiken & Picton, 2008; Aljarboa et al., 2023), mutual information (Cogan & Poeppel, 2011; De Clercq et al., 2023), and regression analysis (Di Liberto et al., 2015). A key advantage to these approaches is that–unlike traditional, event-related paradigms–they allow researchers to employ naturalistic stimuli capable of reflecting the rich spectro-temporal structure and variability present in every-day speech. Such stimuli are likely to engage, and indeed, require, the recruitment of cognitive processes across all levels of speech perception.

Quantifying this form of stimulus-brain coupling over time can shed light on topics as diverse as speech-in-noise perception (McHaney et al., 2021), selective auditory attention (Baltzell et al., 2016), audiovisual integration (Crosse et al., 2015), and infant language acquisition (Barajas et al., 2021). In addition to answering basic science questions about speech perception, there is increasing interest in neural speech tracking as a potential tool for clinicians (Gillis et al., 2022). Such methods have been proposed as objective measures to assess auditory speech processing (Alikovic et al., 2021; Geirnaert et al., 2021; Gillis et al., 2021; Vanthornhout et al., 2018). These techniques provide a means to directly link behaviour with physiology, as the same stimuli and listening paradigms can be used to collect measures from each domain. Still, a detailed understanding of the underlying mechanisms, their reflection in neural response measurements, and potential interactions is required before clinical application. Neural decoding model weights are non-causal and should not be used to draw inferences about the underlying spatial-temporal patterns of brain activity associated with speech perception (Kriegeskorte & Douglas, 2019). They are, however, ideal for clinically-oriented research, as decoders are not computationally constrained, do not require many parameters to be manually selected—such as the pre-identification of channels or sensors of interest—and provide a simple, univariate measure of accuracy. Hence, in the current study, we focus on the neural decoding of speech.

### Interplay Between Sensory and Cognitive Mechanisms During Speech Perception: The Role of Speech Intelligibility

Speech perception emerges at the interface between sensory processing and cognition; that is, the experience of hearing speech results from the active interpretation and representation, rather than deterministic or passive translation, of acoustic signals as higher-order linguistic constructs (Davis & Johnsrude, 2007; Holt & Lotto, 2010; Heald & Nusbaum, 2014). This complexity is reflected by the physiological organisation of the auditory system, which contains numerous and substantial bidirectional connections between cortical and sub-cortical regions (Terrero & Delano, 2015; Asilador & Llano, 2021). In practice, such recurrence in the flow of information across and between levels of the auditory hierarchy suggests that even early sensory processing is affected by computations performed at the cortex (Lesicko & Llano, 2017; Price & Bidelman, 2021). When decoding neural speech tracking, it is therefore possible that contextual factors associated with higher-order processing, such as semantic prediction, may modulate the sensory response to lower-order stimulus features, such as the amplitude envelope. In fact, previous studies reported that neural tracking of acoustic properties was directly affected by manipulations of speech intelligibility, and also significantly correlated with behaviourally measured comprehension in typically hearing adult and children listeners (Iotzov & Parra, 2019; Peelle et al., 2013; Van Hirtum et al., 2023; Vanthornhout et al., 2018), as well as CI listeners (Nogueira & Dolhopiatenko, 2022; Verschueren et al., 2019; Verschueren et al., 2021). Many of these studies did not manipulate speech intelligibility independently, but instead varied speech-to-noise ratios or speech intensity levels as proxies. This approach may lead to changes in physical aspects of the signal, such as reductions in spectral contrast or audibility, which could alter the neural response irrespective of intelligibility *per se*. Instead, another option for manipulating speech intelligibility is to use unfamiliar natural languages. For example, Etard and Reichenbach (2019) presented English-speaking listeners with stimuli in both English and Dutch, reporting significantly greater neural decoding for English when the speech was heard in noise. However, the authors also found that neural decoding was, overall, strongest for speech without noise, in which case, the magnitude of tracking did not differ between English and Dutch (Etard & Reichenbach, 2019). One recent study presented Dutch and Frisian speech to Dutch-speaking listeners and found similar neural decoding in both languages, with the caveat that some participants reported being able to understand the Frisian stimuli (Gillis et al., 2023). These results are further complicated by a separate, but related strand of research that compares speech tracking across the adult language learning process. Such work suggests that the magnitude of speech-neural coupling during second language (L2) listening becomes greater with increasing L2 proficiency (Lizarazu et al., 2021; Zinszer et al., 2022); however, higher L2 neural decoding may also reflect increased attention and/or effortful listening (Reetzke et al., 2021; Song & Iverson, 2018). Indeed, attention and effortful listening may play a role in many studies that manipulate intelligibility: Participants’ capacity and willingness for sustained, engaged listening is likely to depend on whether they can understand the speech or not. For instance, task-directed attention boosts speech-brain coupling specifically at the timescale of complete phrases (Sokoliuk et al., 2021). Simultaneously watching a silent, unrelated cartoon diminishes auditory speech tracking in noise (Vanthornhout et al., 2019). Selective auditory attention paradigms also reveal that the target among competing speakers can be identified based on neural decoding alone (Baltzell et al., 2016; Biesmans et al., 2016; O’Sullivan et al., 2015), due to a proportionately stronger neural response to the focal stimulus (Ding & Simon, 2012; Golumbic et al., 2013; Rimmele et al., 2015). Higher listening effort, in response to degraded or challenging listening conditions, also appears to enhance speech tracking, such as during reverberant speech perception (Fuglsang et al., 2017; Ershaid et al., 2023). In short, successful reconstruction of acoustic features from neural data relies on sensory and cognitive processes originating across brain networks and regions. Hence, the role of speech intelligibility, and its entanglement with attention and listening effort, needs to be considered for neural speech decoding as an objective measure of auditory speech processing (Lesenfants & Francart, 2020).

### Neural Decoding in the Context of Cochlear Implant Listening

One population that may benefit from clinical applications of neural decoding are cochlear implant (CI) recipients. CIs confer a sense of hearing via electrical stimulation of the recipient’s auditory nerve. Over a million CIs have been implanted globally, with patients ranging from infants to the elderly (Zeng, 2022). Typically, CI device fitting is based on subjective patient reports and simple audiometric measures in a time-constrained environment, which may be insufficient for assessing the transmission quality of speech signals. Furthermore, many CI recipients do not experience good speech reception right after switch-on, or have limited verbal communication skills (e.g., infants), impeding their ability to describe the speech transmission quality of their device. After undergoing a time-intensive fitting process in the clinic, it can take weeks or months for an individual to adjust to their CI settings. Unfortunately, it is possible that the CI device was in fact not optimally set up for that patient, and by this point, maladaptive listening strategies may have already become established. Speech benefits should be maximised for every CI recipient, and may crucially depend on the acoustic quality of the perceived speech from day one of CI activation. An efficient and objective measure of naturalistic speech as received by CI recipients would, therefore, present a helpful tool for the device-fitting process in clinics, to assess speech perception outcomes over time, and for the development of improved sound processing strategies in future CI devices. For example, although CIs often can restore speech understanding to a high degree, large inter-patient variability remains (Boisvert et al., 2020; Carlyon & Goehring, 2021). Biophysical factors, such as hearing loss etiology and surgical placement of the CI array, have been shown to explain some of this variability, in addition to complications arising at the electrode-neural interface, such as interactions between stimulating electrodes and auditory nerve fibres (Goehring et al., 2020, 2021). However, it remains unclear why some CI recipients achieve very good speech perception, whereas others do not, and why such large differences in speech reception improvement occur (Holden et al., 2013). Ultimately, the interplay between sensory and cognitive processes discussed in the previous section apply to speech perception with CIs; and given the sparsity and distortions inherent to the perceived CI speech signal, it is likely that central factors are important to speech outcomes in this population (Heydebrand et al., 2007).

Neural speech tracking has been previously measured in CI listeners to examine its relationship with speech reception skills (Verschueren et al., 2019; Verschueren et al., 2021) and to identify the locus of auditory attention (Dolhopiatenko & Nogueira, 2023; Nogueira & Dolhopiatenko, 2022). A major complication when decoding speech from the brain of CI listeners is the electrical stimulation by the CI itself, which is registered by the EEG recording system concurrently with brain-originating activity, thereby confounding the observed neural representation of the speech envelope. To avoid electrical contamination entirely, researchers may instead simulate CI listening in acoustically-hearing listeners using vocoded speech. Vocoder-based CI simulations are inherently limited and cannot account for all aspects of CI hearing (Svirsky et al., 2021), but do enable full experimental control over the perceived speech signal, including its spectral resolution and degradation (Cychosz et al., 2023). A further advantage is that neural responses to CI-simulated speech can be compared to neural responses to unprocessed speech within the same listener as reference.

### Disentangling Intelligibility from Spectral Degradation Effects

Previously, we used vocoding to investigate the consequences of spectral degradation on neural decoding of the speech amplitude envelope in 20 typically hearing listeners using EEG (MacIntyre & Goehring, 2023). Despite relatively severe spectral degradation, we found no statistical evidence for an effect on neural decoding, but only included intelligible speech conditions. Had we introduced acoustically similar, but non-intelligible speech, listeners might not have benefited from cognitive processes related to intelligibility such as semantic prediction. In other words, although we had determined neural tracking of intelligible speech to be robust to spectral degradation, non-intelligible speech tracking could diminish as the signal becomes increasingly spectrally degraded.

The current EEG study is, therefore, motivated to disentangle speech clarity and comprehension as potential factors underlying the success of neural decoding. To achieve this, we independently manipulate speech intelligibility and spectral degradation, taking care that acoustic, linguistic, textual, and speaker-specific differences, which may have been confounded with intelligibility in previous work, are minimised here. For non-intelligible listening conditions, we opted to present our English-speaking participants with natural, Dutch language stimuli (Christophe & Morton, 1998; Ramus et al., 2003). To produce similar English and Dutch speech stimuli, we selected a common text source material and recruited an individual, early bilingual speaker to record in both languages, thereby holding speaker-specific characteristics constant. To manipulate acoustic clarity, in addition to natural (unprocessed) speech, we introduce two levels of spectral degradation with vocoded speech and vocoded speech with simulated current spread as it occurs with CIs (Grange et al., 2017). Note that, to address the issue of waning attention during non-intelligible speech perception, we introduce a prosody-focused auditory target detection task that can be performed with or without speech understanding.

### This Study: Aims and Hypotheses

Our primary aim is to disentangle intelligibility from spectral degradation effects during the neural decoding of speech. To confirm its use for clinical applications, we conduct our EEG analyses using both subject-specific and group-level neural decoding, illuminating what is captured in common by these different approaches. We also investigate the generalisability of decoding by training and testing within and across intelligibility and spectral degradation conditions, similarly to other studies that trained and tested decoders using differing stimulus materials and listening conditions (e.g., Corcoran et al., 2023; Gillis et al., 2021). Finally, it may be clinically important to establish that an individual patient’s neural tracking is decoded significantly above chance, rather than taking the absolute magnitude of decoding accuracy as the objective measure of interest, to account for the substantial variability in neural decoding measures. Here, we employ random permutation testing to describe the proportion of decoders that reach conventional thresholds of statistical significance. Building on the mixed results of the studies most similar to ours, we propose two hypotheses. For the first hypothesis, and in line with reports of an association between comprehension and speech-brain coupling (Iotzov & Parra, 2019; Peelle et al., 2013; Vanthornhout et al., 2018), we predict that speech tracking is generally better for intelligible (English) in comparison to non-intelligible (Dutch) speech. Previous reports suggest that decreasing acoustic quality may in fact enhance neural tracking of intelligible speech (Hauswald et al., 2022; Yasmin et al., 2023); however, as we saw no evidence for a group-level effect of spectral degradation in our previous work (MacIntyre & Goehring, 2023), we do not expect to find one in the current study. In the second hypothesis, we still expect that intelligibility is associated with stronger speech-brain coupling, but only as an interaction effect with spectral degradation. Specifically, whereas English speech tracking remains robust, Dutch speech tracking diminishes, with declining acoustic clarity. This result would cohere with previous works where a language-based difference was only observed for speech in noise (Etard & Reichenbach, 2019) and no difference was found between two languages presented as unprocessed speech only (Gillis et al., 2023).

## Materials and Methods

### Participants

A group of 38 adults (22 female and 16 male; ages 18-35, Mean = 24.95, SD = 5.09; 6 self-reported as left-handed) with self-reported typical hearing took part. The target sample size was determined to roughly double that of previous EEG studies reporting intelligibility effects (e.g., Etard & Reichenbach, 2019; Lesenfants et al., 2019). Participants were recruited from the departmental volunteer database, and comprised university students as well as members of the broader local community. Participants reported no history of audiological nor speech and language-related disorders, and spoke English as a primary language with little to no experience of Dutch. Limited exposure to other related languages, namely Afrikaans and German, was reported by 6 participants, but these individuals verbally confirmed that they could not understand the Dutch stimuli (self-rated “Ability to Follow” Median = 1.75, IQR = 1.00, scale 1–7). All participants provided informed, written consent and received £12.50 per hour for taking part. The study received approval from the Cambridge Psychology Research Ethics Committee and was conducted in accordance with the Declaration of Helsinki.

### Stimuli

#### Recording, Preprocessing, and Editing of Audio

The participants were presented with an audio recording of the Sherlock Holmes story “The Adventure of Charles Augustus Milverton” by Arthur Conan Doyle (Doyle et al., 1903), which is also available as a Dutch translation (Doyle, 1903). Only one of the participants reported having encountered the story previously. For both languages, the text was lightly edited for clarity and modernisation. In Dutch, the names of characters and places were changed (e.g., “Holmes” to “Haas”). An early bilingual, adult male speaker was recruited to perform both versions of the story. The speaker spoke American English at school and Dutch in the home during his childhood. The English story was 33 min. 47 s, divided into 6 trials (Mean = duration 5 min. 38 s, SD = 2.07 s) and the Dutch story was 35 min. 54 s, also divided into 6 trials (Mean = duration 5 min. 59 s, SD = 4.23 s).

Recordings were sampled at 44.1 kHz. Intensity was normalised by root mean square using a sliding window and corrected by ear for a naturalistic narrative experience; the integrated loudness was −19.90 Loudness Unit Full Scale (LUFS) for the English story and −20.03 LUFS for the Dutch version. Audio was filtered to conform to the international long-term average speech spectrum (Byrne et al., 1994; Figure 1, Panel A). Long pauses (*>* 500 ms) were manually restricted to 500 ms. Syllabic timing was estimated by windowing the speech recordings into 2 s chunks and automatically detecting vowel onset-like acoustic landmarks (MacIntyre et al., 2022). The grand average inter-vowel onset interval was, for English, 203.47 ms (SD 47.11 ms) and for Dutch, 202.68 ms (SD 38.96 ms). The grand average coefficient of variation for inter-vowel onset intervals was 0.56 (SD 0.23) for English and 0.53 (SD 0.21) for Dutch. Hence, the English and Dutch speech stimuli were well-matched for overall syllable rate and regularity (Figure 1, Panels C and D).

**Figure 1.**
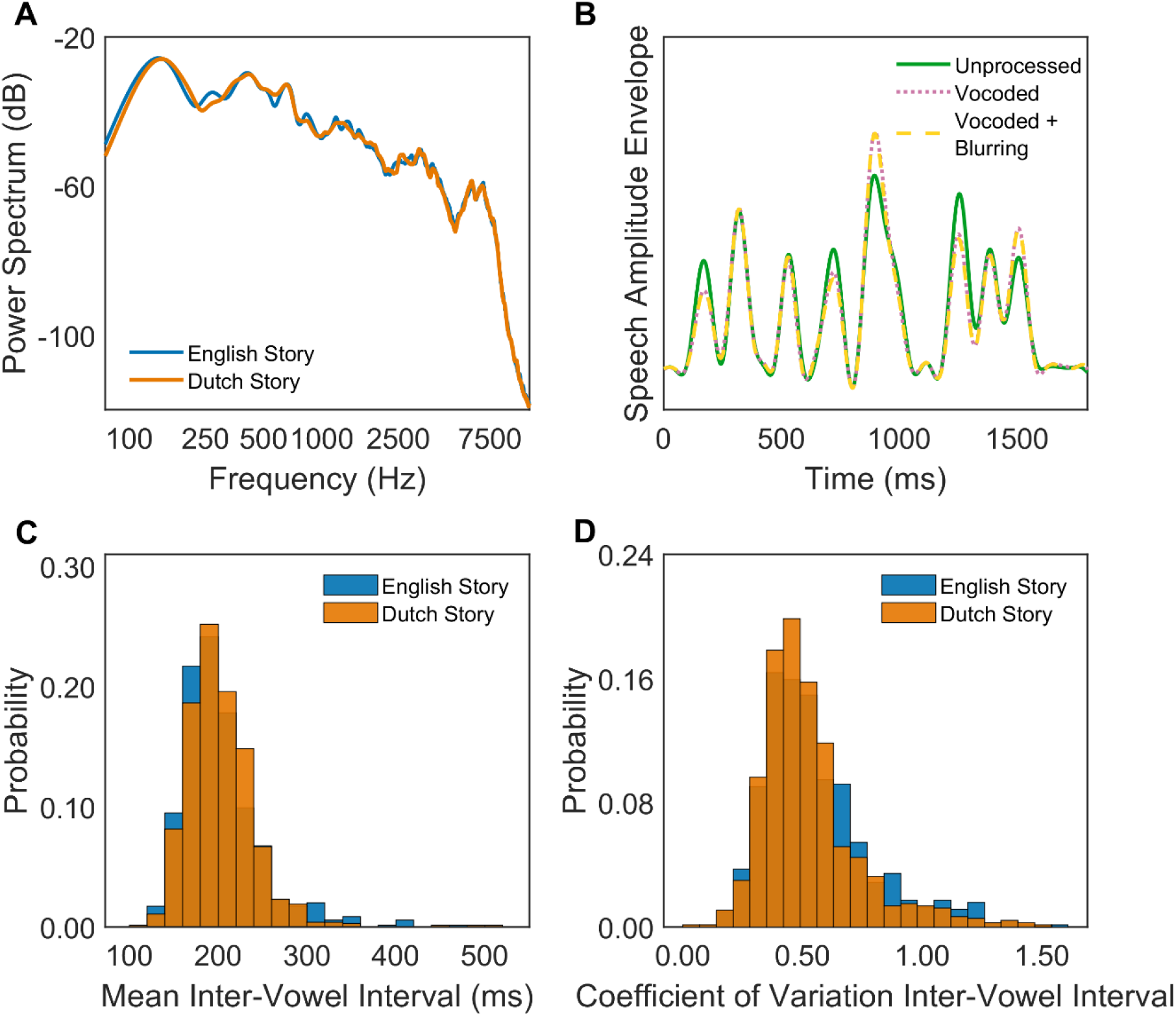
Comparisons of English and Dutch speech stimuli. Panel A: Spectral power analysis of the English and Dutch versions of the story. Panel B: Speech amplitude envelopes extracted from Unprocessed, Vocoded, and Vocoded + Blurring listening conditions. Panel C: Histograms depicting the distributions of mean inter-vowel intervals (calculated over a 2 s-duration window) in the English and Dutch versions of the story. Panel D: Histograms depicting the distributions of coefficient of variation of inter-vowel intervals (calculated over a 2 s-duration window) in the English and Dutch versions of the story.

#### Spectral Degradation Conditions

There were three spectral degradation conditions. The first consisted of natural or unprocessed speech (“Unprocessed”). To produce the second spectral degradation condition (“Vocoded”), we employed the SPIRAL vocoder (Grange et al., 2017). This technique was specifically designed to model CI listening by emulating the physiology of the spiral ganglion and interactions at the electrode-neural interface (Grange et al., 2017). In particular, SPIRAL accounts for the channel interaction mechanism, by which spectral resolution is further degraded with CIs due to overlap in neural excitation, effectively “blurring” the incoming signal (Fu & Nogaki, 2005; Goehring et al., 2020, 2021). Channel interaction also affects temporal speech cues via the smoothing of envelope fluctuations across channels (Oxenham & Kreft, 2014). Behavioural experiments indicate that SPIRAL-based simulations can match typical CI speech reception performance in background noise (Fletcher et al., 2019; Fletcher et al., 2018; Goehring et al., 2019). We used 16 analysis filter bands, resulting in spectrally degraded but intelligible speech. In the third spectral degradation condition (“Vocoded + Blurring”), we additionally simulate the deleterious effects of current spread when vocoding the speech by implementing a decay slope of −16 dB/octave. This value is proposed to best match CI speech reception performance based on behavioural data (Grange et al., 2017). As naive listeners rapidly learn to understand vocoded speech (Hervais-Adelmanet al., 2011; Samuel & Kraljic, 2009), we address perceptual learning effects over the course of the experiment by counter-balancing the 6 combinations of language and spectral degradation condition. Both the English and Dutch stories were presented chronologically (i.e., the narrative structure was preserved), alternating trials between languages. With two trials per listening condition, this resulted in 12 unique condition orders.

### Behavioural Measures

#### Repeated-Phrase Detection Task

The behavioural listening task was designed to sustain auditory attention throughout the experiment, even when the speech stimulus was unintelligible to participants. It consisted of a repeated-phrase detection paradigm. Specifically, the audio would repeat over the course of one short phrase, resulting in one additional occurrence of the phrase after first presentation, after which point the story continued. Such a task avoids the introduction of non-speech target sounds (e.g., tones or noise bursts), or interspersing recognisable English target words within Dutch speech, which may not require active attention to register. Repetitions occurred pseudo-randomly once every ∼45 s and each trial contained 8 repeated phrases in total. The Mean repeated-phrase duration was 2.06 s (SD 0.04 s). This value was determined via online piloting (n = 20 English speakers, ages 18-65), which showed that the task became more difficult for Dutch stimuli as the phrase length exceeded 3 s. Before beginning the experiment, participants were given a practice session with feedback automatically provided on accuracy and reaction times. There were 6 practice trials (duration range 7-9 s), with one trial for each experimental listening condition. Performance in the practice session was monitored by the experimenter to ensure the task was understood. Target detection was recorded via button press using a custom-made USB button box that the participant held in one hand on their lap.

#### Self-Reported Engagement and Ability to Follow

At the end of each trial, participants were asked to rate the preceding story section in response to two questions using a 7-point Likert scale. The questions were “How engaging did you find the story?” and “How well could you follow the story?”, and the order of question presentation alternated by trial. A rating of “1” meant that the participant did not feel at all engaged by the story or that the participant could not follow the story content at all. A rating of “7” meant that the participant felt highly engaged by the story or that they could easily understand the story content. “Engagement” was explained to the participant as feeling actively interested in the plot and/or outcome of the story. “Ability to follow” was explained as comprehension of the plot and dialogue. Participants’ responses were recorded using the left and right arrow keys of a USB keyboard.

#### Preprocessing of Behavioural Data

To quantify performance in the repeated-phrase target detection task, hit and false alarm rates were measured at the trial level. Reaction times were also aggregated at the trial level by taking the median and inter-quartile range. For all behavioural measures (i.e., hit and false alarm rates, median reaction times, and self-report ratings), we took forward the mean value of the two trials associated with each condition for statistical analysis.

### EEG Acquisition and Preprocessing

The experiment was carried out in an acoustically and electromagnetically shielded booth. 64-channel EEG was digitised at 2048 Hz using a BioSemi ActiveTwo EEG system. Scalp electrodes were placed according to the International 10-20 system. Stimuli were presented binaurally via an RME Fireface UCX (RME, Haimhausen, Germany) external soundcard and ER-2 insert earphones (Etymotic Research, Elk Grove Village, Illinois). Presentation levels were held constant across all participants and throughout experimental sessions, with the experimenter verbally confirming with each individual that the stimulus intensity was clearly audible and comfortable. To establish a common time series, EEG triggers were controlled using an additional channel of the soundcard. Participants interacted with the experiment using a custom GUI programmed in Psychtoolbox (Kleiner et al., 2007). While listening to speech stimuli, they were asked to refrain from unnecessary movement and to focus their eyes on a fixation cross (Thaler et al., 2013). Participants had the opportunity to take a short, self-paced break for blinking and stretching after each trial. At the experiment halfway point, the participant was invited to take a longer break along with the offer of a drink and snack. Each experimental session, including EEG preparation and debriefing, lasted approximately 2 − 2.5 hours.

All preprocessing was conducted using custom routines programmed in MATLAB 2020b (MathWorks Inc., Natick, Massachusetts) with functions from the Fieldtrip (Oostenveld et al., 2011) and noisetools (de Cheveigné & Arzounian, 2018) toolboxes. The EEG data were re-referenced to the channel average and resampled at 256 Hz for computational efficiency. Zapline-plus was applied to remove residual line noise (Klug & Kloosterman, 2022). The data were highpass filtered at 0.5 Hz and then lowpass filtered at 40 Hz using 4th order Butterworth filters. Noisy channels were visually identified and excluded. Eye blink artifacts were removed using independent component analysis (runica algorithm). Previously identified noisy channels were then interpolated using a weighted neighbours approach. The first and final 5 seconds of each trial were trimmed to avoid stimulus onset-related responses and filtering-related artifacts. Both the EEG and acoustic envelope signals were resampled to 100 Hz in order to compute the decoders.

### Measure of Cortical Speech Tracking

The acoustic stimulus envelope was extracted by passing the speech signal through a Gammatone filter-bank (28 filters equally spaced on the equivalent rectangular bandwidth scale between 50 and 5000 Hz); rectifying and applying a power law to each sub-band output; and then recombining the resultant envelopes by taking their average (Biesmans et al., 2016). Finally, the envelope was lowpass filtered at 10 Hz using an 8th order Butterworth filter to form the broadband envelope. A linear comparison of envelopes across the three levels of spectral degradation revealed that all pairwise correlations were very strong (*r >* 0.985, *p <* 0.001).

To assess neural tracking of the speech envelope, we trained linear decoders using the Multivariate Temporal Response Function (mTRF) Toolbox (Crosse et al., 2016). The mTRF Toolbox applies regularised ridge regression to obtain a quantitative mapping between stimulus features–here, the speech amplitude envelope–and the recorded neural response. The objective is to reconstruct or predict the input stimulus from EEG (Crosse et al., 2016, 2021). Within this framework, the decoding model *g* (*τ, n*) returns the stimulus *s*^(*t*) as a linear convolution of the neural response *r* (*t, n*) over a range of time lags *τ*, or

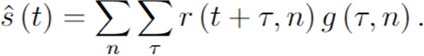

The accuracy of the linear decoder can be estimated by finding the correlation (in this case, Pearson’s *r*) between the reconstructed and true input features. This measure of neural speech tracking formed the primary dependent variable in our study. Data to be held out for prediction were selected as the first 60 s of the later of the two trials associated with each listening condition.

When considering practical applications for example in a clinical context, it is currently unknown whether it would be preferable to build decoder models on a patient-by-patient basis or whether a generic decoder model should be used for assessing speech envelope decoding outcomes, and whether these two approaches would produce different results. We therefore trained two types of decoders: subject-specific models using data from individual participants and group-level models using data pooled across participants. We trained the decoders by minimising the mean squared error using *k*-fold, leave-one-out cross-validation (Crosse et al., 2016, 2021). In the case of the participant-specific decoders, we split the ∼10 minutes of training data into 4 folds; for the group decoders, we trained on *n*-1 × ∼10 minute folds. The time lags encompassed the range from −50 to 400 ms post-stimulus. The optimised ridge parameter *λ*, which controls the regularisation penalty, was empirically determined using the same format of *k*-fold cross-validation within logarithmically spaced values ranging between 10^−7^ − 10^7^. As our goal was to maximise decoding accuracy with a view to clinical applications, an individual *λ* value was chosen for each model; however, optimal *λ* was 100 for the majority (30/38) of subject-specific decoders (Range = [10, 1000]), and 1000 for all group-level decoders.

### Statistical Analysis

To estimate the effects of intelligibility and spectral degradation on speech envelope reconstruction accuracy whilst accounting for differences across participants, we conducted linear mixed effect modelling in R 4.0.3 (R Core Team, 2021) with lme4 (Bates et al., 2014). These models were estimated using restricted maximum likelihood and optimx optimizer. When modelling neural decoding accuracy as an *r*-value, the primary predictors of interest were the factors Language (Trained and Test) and Spectral Degradation (Trained and Test). Random intercepts and slopes within Participant were fit subject to model convergence. The reference levels were Trained and Test on English, Unprocessed speech. Model residuals were inspected using visualisations and diagnostic tests of normality from the DHARMa package (Hartig, 2017). The selection of fixed effect terms was guided by comparing models with the Akaike information criterion (AIC), a metric that balances the model’s goodness of fit against its parsimony (Burnham et al., 2011). The significance of fixed effects was tested using Type II Wald *F* tests with Kenward-Roger degrees of freedom (Luke, 2017) using the pbkrtest package (Halekoh & Højsgaard, 2014), and *R*^2^ values (calculated with r2glmm package) are provided as a measure of variance explained by each predictor (Jaeger et al., 2017). Planned and posthoc comparisons were conducted using estimated marginal means implemented in easystats package (Lüdecke et al., 2022). For correlational analyses, continuous variables of interest included Hit Rate, False Alarm Rate, and Mean Median Reaction Time in the repeated-phrase detection task, as well as the ordinal variables of self-reported Engagement and Ability to Follow.

Bonferroni-adjusted *p*-values are reported where relevant. In-text confidence intervals are obtained by using bootstrapped sampling with replacement (*n* = 1000 iterations).

#### Establishing Decoder Significance

The primary outcome variable is decoder accuracy as an *r*-value, following convention. However, the individual significance level of each decoder, relative to an empirically determined null distribution, may also be informative. To obtain decoder significance, we first generated a null distribution by segmenting the training speech envelope into ∼3 s chunks, shuffling the segments (prohibiting any coincidental true pairings), and re-concatenating them. Segmentation was applied at signal zero-crossings and the segments were smoothly joined using interpolation, thereby avoiding the introduction of artefacts to the permuted envelope. We then trained and tested null decoders, within-listening condition, using this shuffled stimulus and the unmodified neural response 1000 times. When the observed *r*-value exceeds the 95^th^ percentile of the null distribution, the decoder can be considered as significant at *p <* 0.05. We describe the frequency and variability of decoder significance by listening condition.

## Results

### Behavioural Measures

#### Self-Reported Ability to Follow and Engagement

Summary descriptive statistics for the behavioural results by listening condition are shown in Table 1 and Figure 2. Averaging across levels of acoustic clarity, participants rated their Ability to Follow the English story as Median = 6.00 (IQR = 2.0), and the Dutch story, Median = 1.00 (IQR = 0.50) on a scale from 1 − 7. Self-reported Engagement was high in English, with even the least engaging listening condition, Vocoded + Blurring, receiving a median score of 5.50 (IQR = 3.00). For Dutch, however, the overall median Engagement score was just 1.50 (IQR = 1.00) on a scale from 1 − 7.

**Figure 2.**
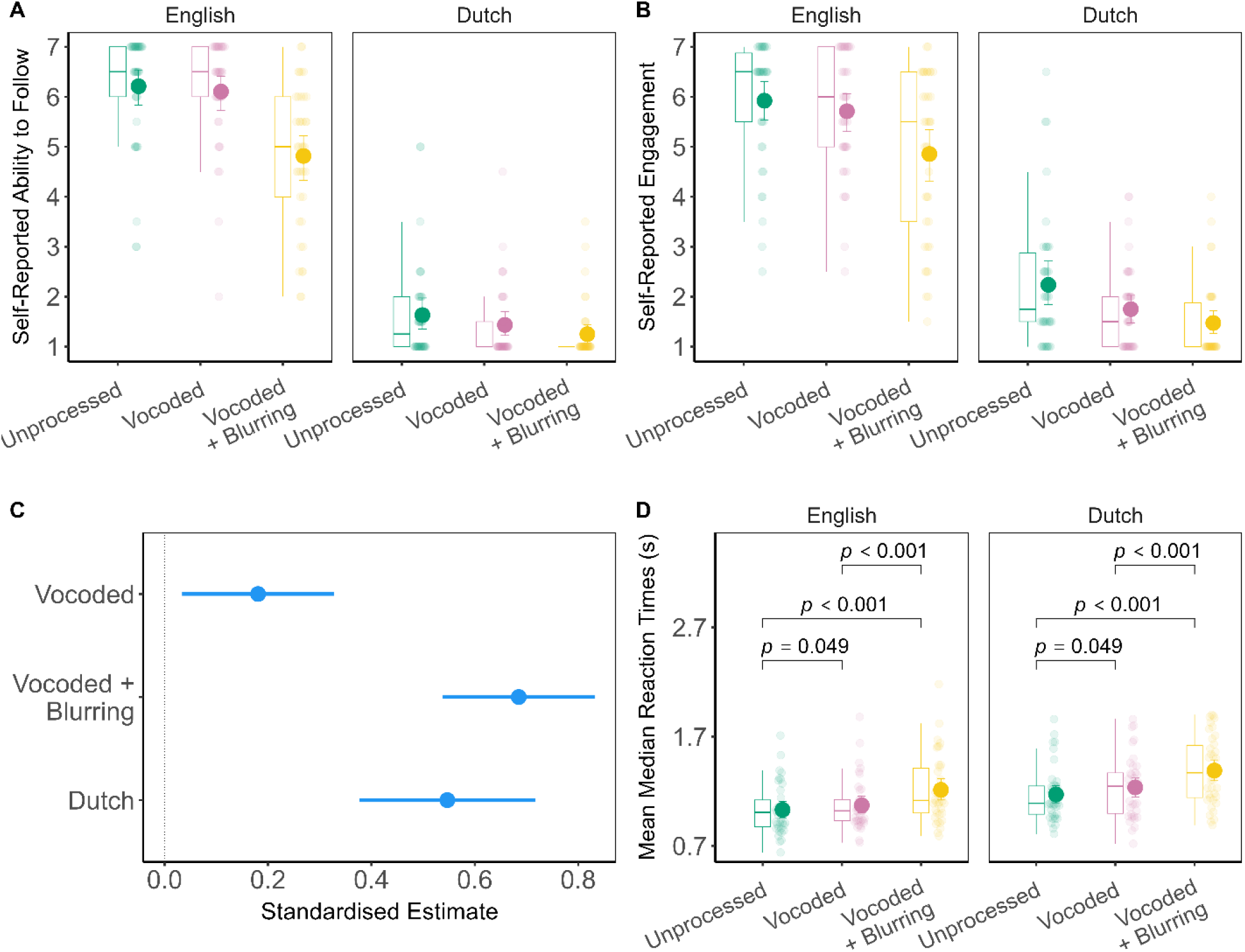
Panel A: Box plots and group mean with bootstrap 95% confidence intervals depicting self-reported ability to follow the story. Panel B: Box plots and group mean with bootstrap 95% confidence intervals depicting self-reported engagement with the story. Panel C: Dot-and-whisker plot depicting standardised estimates (regression coefficients) with 95% confidence intervals from the linear mixed effects model of mean median reaction times in the repeated-phrase target detection task. The model reference levels were English Unprocessed. Panel D: Box plots and group mean with bootstrap 95% confidence intervals depicting mean median reaction time by listening condition. Significance values reflect the outcome of pairwise tests of estimated marginal means between levels of spectral degradation with Bonferroni correction.

**Table 1.**
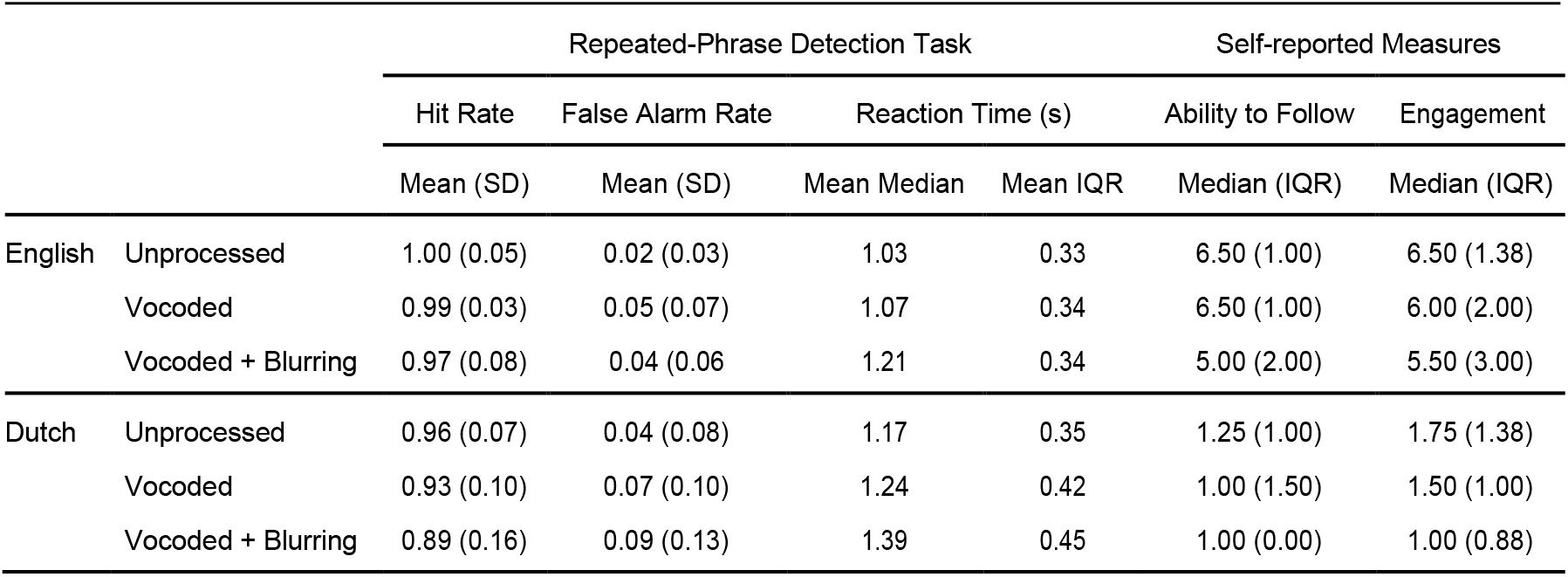
Summary statistics of behavioural data for the repeated-phrase detection task and the participants’ self-reported measures of ability to follow the story in terms of plot and dialogue, and engagement, referring to their sense of absorption or interest in the outcome of the story.

|  |  | Repeated-Phrase Detection Task |  |  |  | Self-reported Measures |  |
| --- | --- | --- | --- | --- | --- | --- | --- |
|  |  | Hit Rate | False Alarm Rate | Reaction Time (s) |  | Ability to Follow | Engagement |
|  |  | Mean (SD) | Mean (SD) | Mean Median | Mean IQR | Median (IQR) | Median (IQR) |
| English | Unprocessed | 1.00 (0.05) | 0.02 (0.03) | 1.03 | 0.33 | 6.50 (1.00) | 6.50 (1.38) |
|  | Vocoded | 0.99 (0.03) | 0.05 (0.07) | 1.07 | 0.34 | 6.50 (1.00) | 6.00 (2.00) |
|  | Vocoded + Blurring | 0.97 (0.08) | 0.04 (0.06) | 1.21 | 0.34 | 5.00 (2.00) | 5.50 (3.00) |
| Dutch | Unprocessed | 0.96 (0.07) | 0.04 (0.08) | 1.17 | 0.35 | 1.25 (1.00) | 1.75 (1.38) |
|  | Vocoded | 0.93 (0.10) | 0.07 (0.10) | 1.24 | 0.42 | 1.00 (1.50) | 1.50 (1.00) |
|  | Vocoded + Blurring | 0.89 (0.16) | 0.09 (0.13) | 1.39 | 0.45 | 1.00 (0.00) | 1.00 (0.88) |

#### Repeated-Phrase Detection Task

Accuracy in auditory target detection ranged from a Mean Hit Rate of 0.89 (SD 0.16) and a Mean False Alarm Rate of 0.09 (SD 0.13) for Dutch Vocoded + Blurring, to a Mean Hit Rate of 1.00 (SD 0.05) and a Mean False Alarm Rate of 0.02 (SD 0.03) for English Unprocessed. Hence, task performance was generally good across all listening conditions. The linear mixed effect model analysis of reaction time data (Figure 2, Panel C) consisted of a main effect of Language, *F*(1.00,37.00) = 42.25, *p <* 0.001, *R*^2^ = 0.53, 95% CI [0.34,0.71], and of Spectral Degradation, *F*(2.00,150.00) = 45.58, *p <* 0.001, *R*^2^ = 0.38, 95% CI [0.27,0.49]. Including an interaction term between Language and Spectral Degradation did not improve model fit (Appendix A, Table A1).

Reaction times to Dutch targets (Mean = 1.26 s, SD = 0.29 s) were slower than to English targets (Mean = 1.10 s, SD = 0.27 s), Estimate = 0.16, 95% CI [0.11, 0.21], *t*(37.00) = 6.50, *p<* 0.001. With respect to Spectral Degradation, in comparison to Unprocessed speech (Mean = 1.10 s, SD = 0.25 s), Vocoded speech (Mean = 1.15 s, SD = 0.27 s) evoked slower reaction times, Estimate = 0.05, 95% CI [0.01, 0.10], *t*(150.00) = 2.43, *p* = 0.016. Vocoded + Blurring (Mean = 1.30 s, SD = 0.32 s) was also associated with slower reaction times in comparison to Unprocessed speech, Estimate = 0.20, 95% CI [0.16, 0.24], *t*(150.00) = 9.21, *p <* 0.001. Finally, Vocoded + Blurring reaction times were also slower than those evoked by Vocoded speech, Estimate = −0.15, 95% CI [−0.20, −0.09], *t*(150.00) = −6.78, *p_bonf_ <* 0.001. Standardised estimates and additional model details are provided in Appendix A, Tables A2−A3).

In summary, reaction times to Dutch and spectrally degraded repeated-phrase targets tend to be slower than responses to English Unprocessed targets. There does not appear to be an interaction between Language and Spectral Degradation, however. Linking response speed to the self-report measures, we found that participants with faster reaction times to auditory targets on average tended to have worse self-described Ability to Follow, *r*_s_ = −0.58, 95% CI [−0.73, −0.17], *p <* 0.001; as well as lower levels of Engagement, *r*_s_ = −0.43, 95% CI [−0.67, −0.21], *p* = 0.007.

### Neural Decoding of the Speech Signal

#### Subject-specific Decoding Accuracy

Each subject-specific decoder was trained on 10 minutes of data from a single listening condition, and tested on 1 minute of held-out test data from each of the six listening conditions. The linear mixed effect model of subject-specific decoding accuracy (Figure 3) included main effects of Trained: Spectral Degradation, *F*(2.00,1245.00) = 9.42, *p <* 0.001, *R*^2^ = 0.01, 95% CI [0.01,0.03], and Test: Spectral Degradation, *F*(2.00,37.00) = 4.16, *p* = 0.023, *R*^2^ = 0.18, 95% CI [0.04,0.44], as well as their interaction, *F*(4.00,1245.00) = 41.74, *p <* 0.001, *R*^2^ = 0.12, 95% CI [0.09,0.15]. In addition, the model also included a non-significant predictor of Trained: Language, *F*(1.00,1245.00) = 3.45, *p* = 0.064, *R*^2^ = 0.00, 95% CI [0.00,0.01], and a main effect of Test: Language, *F*(1.00,1245.00) = 55.29, *p <* 0.001, *R*^2^ = 0.04, 95% CI [0.02,0.07], as well as an interaction between Trained: Language and Test, *F*(1.00,1245.00) = 26.34, *p <* 0.001, *R*^2^ = 0.02, 95% CI [0.01,0.04]. AIC indicated that including interaction terms between the Spectral Degradation- and Language-based predictors did not improve model fit (Appendix A, Table A4). Full details are reported in Appendix A, Table A5.

**Figure 3.**
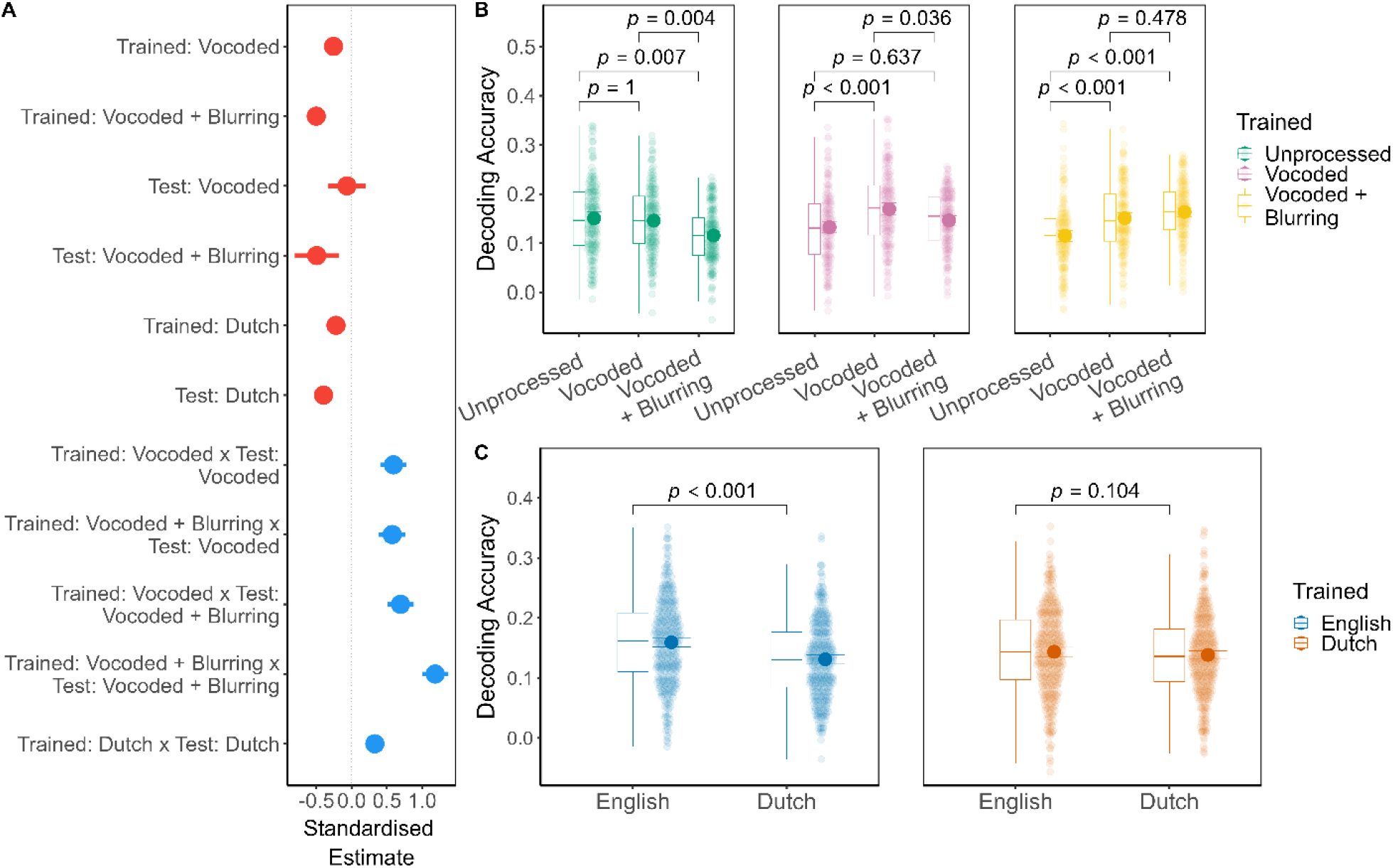
Subject-specific speech decoding accuracy. Panel A: Dot-and-whisker plot depicting standardised estimates (regression coefficients) with 95% confidence intervals from the linear mixed effects model of subject-specific speech decoding accuracy. The model reference levels were English Unprocessed (Trained and Test), with red colours depicting negative estimates, and blue, positive estimates. Panel B: Box plots and group mean with bootstrap 95% confidence intervals depicting speech decoding accuracy by Trained: Spectral Degradation and Test: Spectral Degradation.Data are collapsed across Language conditions.Panel C: Box plots and group mean with bootstrap 95% confidence intervals depicting speech decoding accuracy by Trained: Language and Test: Language.Data are collapsed across Spectral Degradation conditions. Significance values reflect the outcome of pairwise tests of estimated marginal means with Bonferroni correction.

##### Spectral Degradation

To investigate the interaction term between Trained: Spectral Degradation and Test: Spectral Degradation, we conducted pairwise comparisons of the estimated marginal means (Figure 3, Panel B; Appendix A, Table A6). In general, the more similar Trained: Spectral Degradation was to Test: Spectral Degradation, the more decoding accuracy improved. For example, if the decoder was Trained: Unprocessed, then Test: Unprocessed (Mean = 0.15, SD = 0.08) was better decoded than Vocoded + Blurring (Mean = 0.12, SD = 0.07), *p_bonf_* = 0.007, *d* = 0.46, 95% CI [0.16, 0.75]. An analogous, but reverse pattern was observed for Trained: Vocoded + Blurring decoders, such that Test: Vocoded + Blurring (Mean = 0.16, SD = 0.06) was decoded more accurately than Unprocessed (Mean = 0.12, SD = 0.06), *p_bonf_ <* 0.001, *d* = −0.64, 95% CI [−0.94, −0.33]. Decoders that were Trained: Unprocessed or Vocoded + Blurring could each decode Test: Vocoded speech as well as the listening condition they had been trained on (*p_bonf_* ≥ 0.478). However, when the decoder was itself Trained: Vocoded, then Test: Vocoded elicited higher decoding accuracy (Mean = 0.17, SD = 0.08) than both Unprocessed (Mean = 0.15, SD = 0.07), *p_bonf_ <* 0.001, *d* = −0.54, 95% CI [−0.82, −0.25]; as well as Vocoded + Blurring (Mean = 0.15, SD = 0.07), *p_bonf_* = 0.036, *d* = 0.35, 95% CI [0.08, 0.61].

##### Language

We next examined the interaction between Trained: Language and Test: Language (Figure 3, Panel C; Appendix A, Table A7). When Trained: English, decoding accuracy was higher for Test: English (Mean = 0.16, SD = 0.07) in comparison to Dutch (Mean = 0.13, SD = 0.06), *p_bonf_ <* 0.001, *d* = 0.25, 95% CI [0.20, 0.31]. Decoders that were Trained: Dutch, however, did not distinguish between Test: English (Mean = 0.14, SD = 0.07) or Dutch (Mean = 0.14, SD = 0.07), *p_bonf_* = 0.104, *d* = 0.05, 95% CI [−0.01, 0.10]. Thus, contrary to English, there was no apparent advantage for testing on Dutch when having trained on Dutch. Although the effect of Test: Language was modest, it was relatively consistent across participants.

##### Correlations between Brain and Behaviour

In the following analysis, Trained and Test listening conditions were always matched. As group performance in the repeated-phrase detection task was close to ceiling, it was impractical to correlate Hit or False Alarm Rate with neural decoding accuracy. The only listening condition with some variance in target detection success was Dutch Vocoded + Blurring; however, we observed no trend linking speech tracking to Hit Rate, *p* = 0.616, nor False Alarm Rate, *p* = 0.532. With regards to reaction times, slower response speed was associated with better neural decoding accuracy in English Unprocessed, *r* = 0.38, 95% CI [0.09, 0.60], *p* = 0.019, but not for any other listening condition (*p* ≥ 0.242). The wide confidence intervals of the estimate suggest that, if truly present, this correlation would be subject to inter-individual variability. Turning to self-reported Ability to Follow, we correlated this measure with neural decoding in English Vocoded + Blurring–the only intelligible condition with between-participant differences–and found no obvious association, *p* = 0.504. However, participants differed in their use of the Likert scale, with some participants using its full range across experimental conditions, and others not. To account for this, we subtracted Ability to Follow in English Vocoded + Blurring from English Unprocessed. This gives an individualised measure of relative change in comprehension. We also performed the same operation for decoding accuracy, to produce a measure of relative change in speech tracking. At the group level, there was a correlation between change in Ability to Follow and change in neural tracking, *r* = 0.35, 95% CI [0.02, 0.60], *p* = 0.030. In other words, participants who rated their Ability to Follow as relatively poor for English Vocoded + Blurring were also more likely to have relatively worse speech tracking in that condition, in comparison to English Unprocessed.

Whereas this result coheres with proposals that EEG can predict intelligibility (e.g., Iotzov & Parra, 2019; Vanthornhout et al., 2018), we cannot be sure that changes in decoding accuracy are solely attributable to a loss in intelligibility and were not influenced by acoustic differences. To tease apart these factors, we took advantage of the Dutch decoding data, by subtracting neural decoding in Dutch Vocoded + Blurring from Unprocessed. This third variable provides an independent estimate of individual response to Spectral Degradation that is not expected to vary with intelligibility. We used it as a covariate to calculate the partial correlation between change in Ability to Follow and change in neural tracking in English, showing that *r* = 0.29, 95% CI [−0.11, 0.60], *p* = 0.077. Thus, the association between self-reported comprehension and neural decoding does appear to be mediated by Spectral Degradation, although the wide confidence intervals of the effect suggest considerable inter-individual variability. As Engagement closely tracked with Ability to Follow, we did not examine it further.

#### Group Decoding

Group-level decoding was performed similarly to subject-specific decoding, except instead of training the decoder on a single participant, we used data from *n* −1 participants and held out the *n*^th^ participant’s data for testing. This procedure was repeated for each of the 38 participants. The linear mixed effects model (Figure 4) of group decoding accuracy contained a main effect of Trained: Spectral Degradation, *F*(2.00,1247.00) = 7.07, *p* < 0.001, *R*^2^ = 0.01, 95% CI [0.00,0.03], and non-significant predictor of Test: Spectral Degradation, *F*(2.00,37.00) = 2.21, *p* = 0.124, *R*^2^ = 0.11, 95% CI [0.01,0.35]; in addition, there was an interaction between Trained: Spectral Degradation and Test: Spectral Degradation, *F*(4.00,1247.00) = 17.70, *p <* 0.001, *R*^2^ = 0.05, 95% CI [0.03,0.08]. Unlike the subject-specific decoders, AIC indicated that the predictor Trained: Language did not improve model fit, neither as a main effect nor interaction term (Appendix A, Table A8). There was, however, a main effect of Test: Language, *F*(1.00,1247.00) = 49.19, *p <* 0.001, *R*^2^ = 0.04, 95% CI [0.02,0.06]. Full details are provided in Appendix A, Table A9.

**Figure 4.**
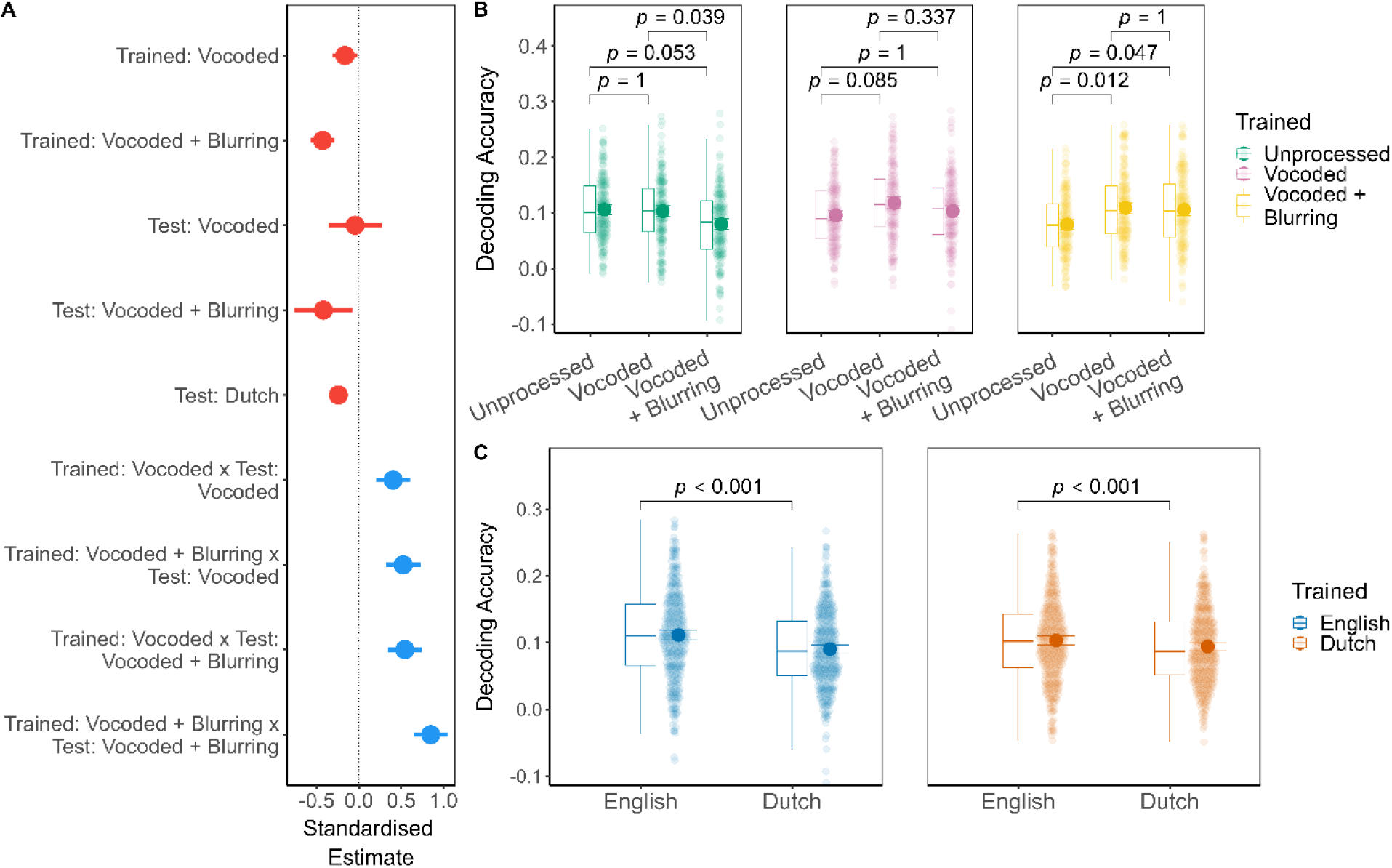
Same as Figure 3 but for group speech decoding accuracy.

##### Spectral Degradation

The pattern of the interaction between Trained: Spectral Degradation and Test: Spectral Degradation was similar to subject-specific decoding. However, negative effects of mismatching Trained and Test conditions were less consistent (Figure 4, Panel B). For example, when the decoder was Trained: Unprocessed, the statistical effect of Test: Unprocessed (Mean = 0.11, SD = 0.06) was only marginally better than Vocoded + Blurring (Mean = 0.08, SD = 0.06), *p_bonf_* = 0.053, *d* = 0.35, 95% CI [0.06, 0.64]. In addition, when the decoder was Trained: Vocoded + Blurring, then Test: Vocoded + Blurring (Mean = 0.11, SD = 0.06) was decoded with only-just significantly higher accuracy than Unprocessed (Mean = 0.08, SD = 0.06), *p_bonf_* = 0.047, *d* = −0.36, 95% CI [−0.65, −0.07]. Finally, at the group level, Trained: Vocoded decoders achieved comparable reconstruction whether Test: Unprocessed, Vocoded, or Vocoded + Blurring. Importantly, for all pairwise contrasts, the wide intervals of the estimates and effect sizes indicate substantial variability in the neural response to Spectral Degradation between participants. Full details of the tests of estimated marginal means are given in Appendix A, Table A10.

##### Language

As indicated by the main effect, Test: English (Mean = 0.11, SD = 0.07) was decoded with higher accuracy than Dutch (Mean = 0.09, SD = 0.06), Estimate = 0.02, 95% CI [0.01, 0.02], *t*(1247.00) = 7.01, *p_bonf_ <* 0.001, Cohen’s *d* = 0.20, 95% CI [0.14, 0.25], regardless of the training language. As with the subject-specific decoding, the Test: Language effect was small in magnitude, but more consistent across participants when compared to the effects of Spectral Degradation.

##### Correlations between Brain and Behaviour

Similarly to the subject-specific results, in the Dutch Vocoded + Blurring listening condition, there was no clear linear relationship between speech tracking and Hit Rate (*p* = 0.154) nor False Alarm Rate (*p* = 0.109). The positive association with slower reaction times in English Unprocessed was present, but numerically smaller than for subject-specific decoding, *r* = 0.33, 95% CI [0.01, 0.04], *p* = 0.041; all other *p* ≥ 0.063). Unlike the subject-specific analysis, however, change in self-reported Ability to Follow and change in decoding accuracy did not significantly correlate, *r* = 0.26, 95% CI [−0.01, 0.64], *p* = 0.121.

#### Generalisability of Neural Decoding

##### Subject-Specific and Group Decoding

To quantify the similarity between subject-specific and group-level decoding performance across listening conditions, we calculated Kendall’s concordance coefficient, *W*, within participant. This was performed using data where the decoder was trained and tested within the same listening condition. For 30 of 38 participants, agreement between their subject-specific and group-level decoding results was *W >* 0.70. The median value was 0.79 (IQR = 0.20, Range = [0.29, 0.97]), which indicates good to very good agreement between subject-specific and group-level decoding accuracy within participant. Although Kendall’s *W* shows high consistency in terms of the direction or ranking of results across listening condition, we were also interested in absolute differences in accuracy between subject-specific and group-level decoding. We, therefore, subtracted each group decoder’s accuracy from each subject-specific decoder’s accuracy, within participant and by listening condition, before aggregating at the participant level. The group mean difference–a loss–in accuracy was 0.06 (SD = 0.04). We can scale this difference by dividing it by the subject-specific decoding value (within participant), resulting in the group Mean = 0.31, SD = 0.20. This can also be described as an average reduction in decoding accuracy of 30% from the subject-specific value (Figure 5, top panel).

**Figure 5.**
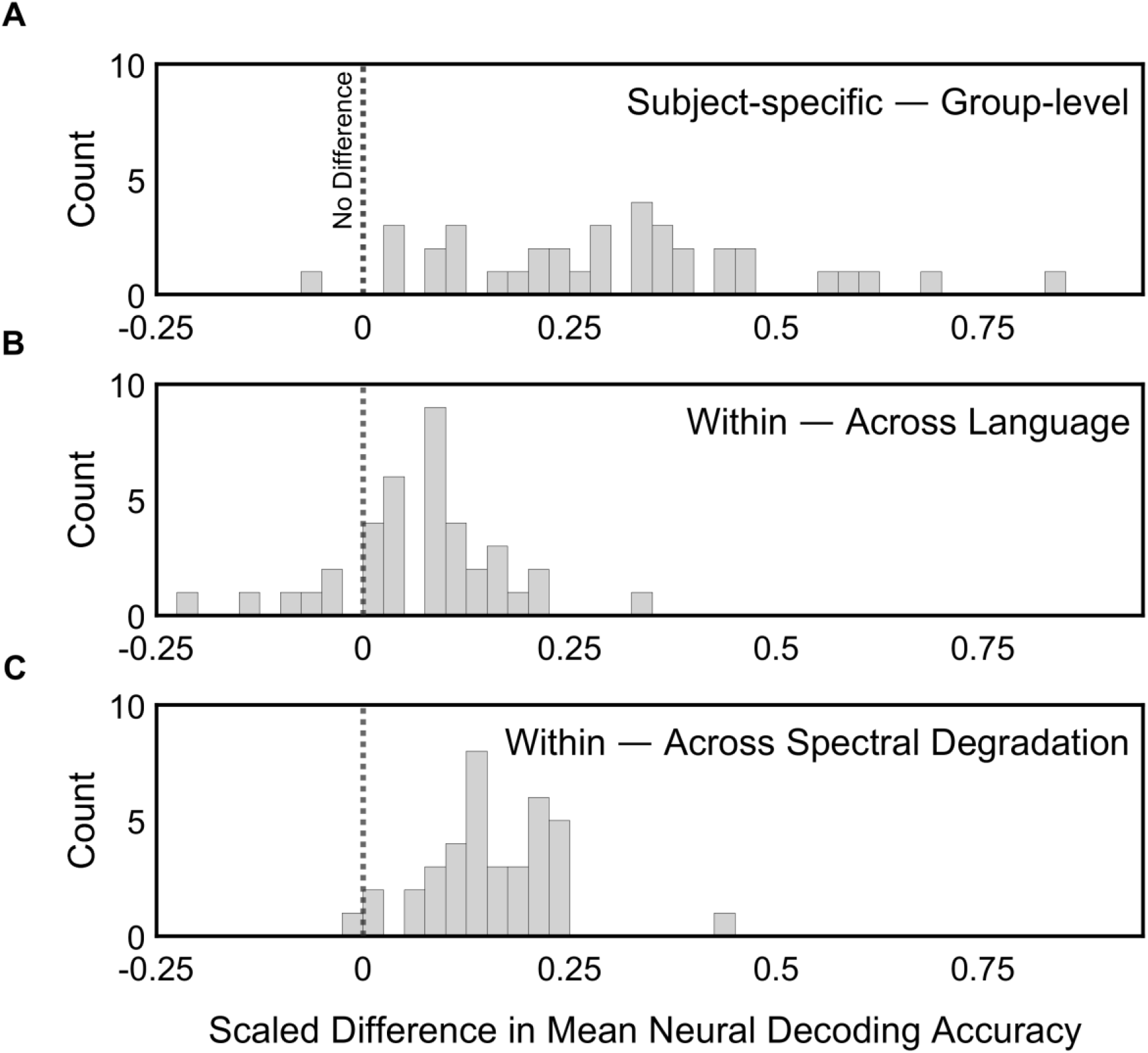
Histograms depicting the scaled difference in neural decoding accuracy when subtracting group-level from subject-specific values (Panel A), or when subtracting non-matched from matched training and testing listening conditions (Panels B and C). Scaling is performed by dividing the resulting differences by the subject-specific (Panel A) or matched training and testing (Panels B and C) decoding accuracy values.

##### Testing and Training Across Listening Conditions

We also examined the global effects of Language and Spectral Degradation when Trained and Test listening conditions are mismatched. Namely, within participant, we subtracted the mismatched (“across”) decoding value from the matched (“within”) decoding value. First, to examine the Language contrast, we pooled subject-specific and group decoding, as well as all three levels of Spectral Degradation. The mean loss in accuracy due to training and testing across Language was 0.01 (SD = 0.01), or when scaled by the within-Language accuracy, Mean = 0.07, SD = 0.10 (Figure 5, middle panel). Next, to look at the Spectral Degradation contrast, we pooled subject-specific and group decoding, as well as Language. In this case, the mean loss in accuracy was 0.02 (SD = 0.01), or when scaled, Mean = 0.16, SD = 0.08 (Figure 5, bottom panel). A paired t-test of the scaled differences confirmed that the proportionate effects of training and testing across listening condition are greater for Spectral Degradation than for Language, *t*(37) = −5.84, *p <* 0.001, *d* = 0.96, 95% CI [−1.34, −0.57].

#### Significance of Decoders

Under clinical settings, it may be useful to select a cut-off value at which point a patient’s neural tracking can be said to be statistically significant. We, therefore, performed a descriptive analysis of decoder significance with a focus on outcomes for individual participants, rather than group-level effects. Decoder statistical significance was established by comparing observed accuracy (i.e., the *r*-value) ranked against the accuracy of decoders trained on randomly permuted versions of the same stimulus speech amplitude envelopes (*n* = 1000 permutations). To quantify the variability of the percentile rank value itself, we calculated median absolute deviation (MAD) at the group level. MAD is a measure of spread analogous to standard deviation, but is more robust to outliers and appropriate for characterising non-Gaussian distributions. Hence, we use MAD here to estimate the consistency, across participants, of the decoder percentile rank relative to its empirically determined, null distribution.

##### Subject-Specific Decoding

Figure 6, Panel A depicts the proportion of subject-specific decoders that met typically accepted levels of statistical significance. English Unprocessed speech was the only listening condition where subject-specific decoding at *p <* 0.05 occurred for every participant. Moreover, MAD = 0.65 percentiles (95% CI [0.36, 1.14]), indicating little variability in the rank of observed decoder accuracy across participants. For English Vocoded speech, however, MAD = 5.05 percentiles (95% CI [0.93, 12.99]); and for English Vocoded + Blurring, MAD = 5.94 percentiles (95% CI [2.08, 14.01]). These larger values of MAD and wider confidence intervals may indicate that, in the current sample, increasing Spectral Degradation resulted in less consistent significance testing: In particular, for English, the proportion of decoders significant at *p* < 0.001 grew from 58% in Unprocessed speech to 71% for Vocoded + Blurring, but so did the proportion of decoders that failed to reach significance at *p <* 0.05, from 0% to 11%, respectively. Turning to Dutch Unprocessed, 21% of decoders (*n* = 8) were non-significant at *p* < 0.05 in the current sample. The percentile rank of Dutch Unprocessed accuracy was also estimated to be the most variable across all the listening conditions, MAD = 9.67 percentiles (95% CI [4.65, 16.43]). Moreover, opposite to the pattern observed for English decoding, variability in significance testing decreased for Dutch Vocoded, MAD = 3.93 percentiles (95% CI [1.44, 10.70]) and Dutch Vocoded + Blurring, MAD = 1.34 percentiles (95% CI [0.50, 2.77]).

##### Group Decoding

The proportions of group decoders to achieve conventional levels of significance were roughly similar across both English and Dutch, with ≥ 28% of decoders failing to surpass the *p <* 0.05 cut-off across listening conditions (Figure 6, Panel B). In other words, about one third of group decoders may not meet a clinically acceptable threshold of statistical significance in the current sample. With respect to the variability of percentile rank across decoders, MAD was smallest, or least dispersed, for Unprocessed speech, with roughly overlapping confidence intervals across English, MAD = 6.06, (95% CI [3.20, 12.05]) and Dutch, MAD = 6.81 (95% CI [4.16, 11.63]). This level of variability was also comparable for Dutch Vocoded, MAD = 7.44 (95% CI [3.52, 12.44]) and Dutch Vocoded + Blurring (MAD = 8.08, 95% CI [5.12, 13.23]). We observed possibly higher MAD with wider confidence intervals for English Vocoded, MAD = 10.52 (95% CI [5.23, 18.47]), as well as English Vocoded + Blurring (MAD = 12.12, 95% CI [6.20, 21.49]). Hence, like the subject-specific decoding, we find numerical differences in the frequency and variability of decoding significance levels achieved by listening condition. However, unlike the subject-specific decoding, the broad pattern of differences are shared across English and Dutch speech.

**Figure 6.**
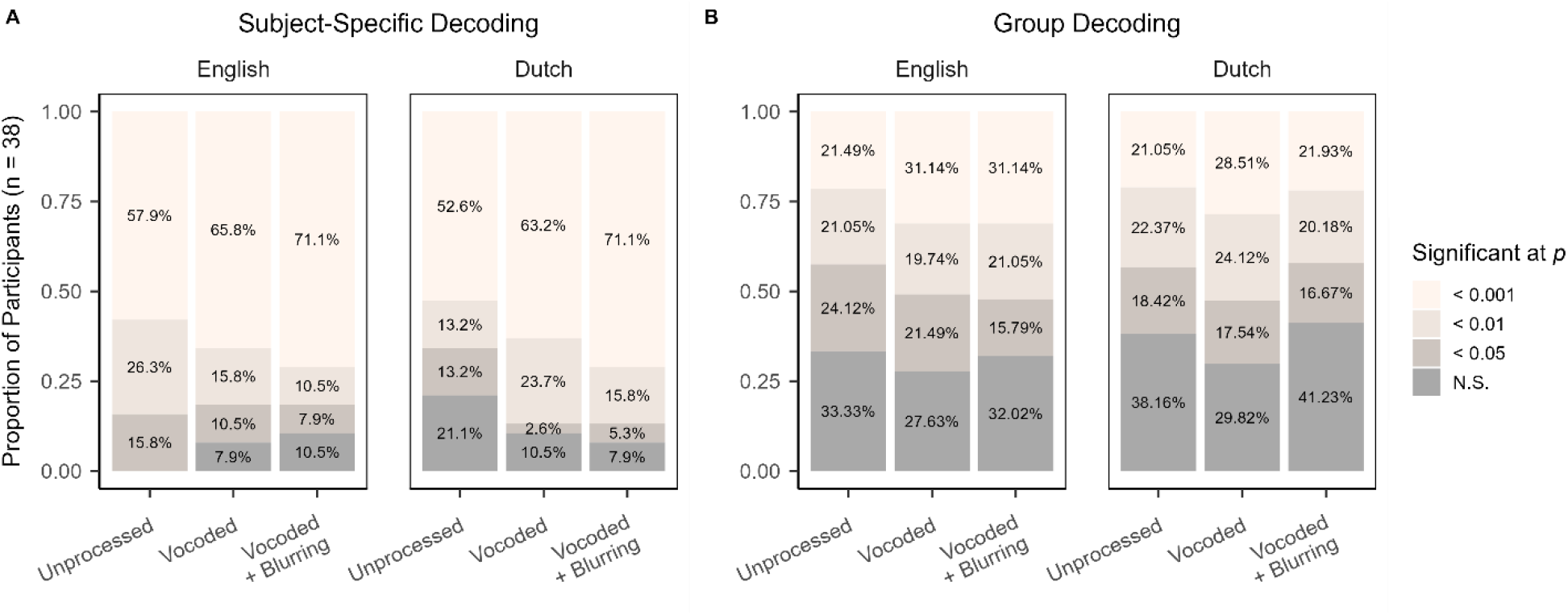
Bar plots indicating the significance level, determined via random permutation testing with n = 1000 iterations, at which neural decoding was decoded across listening conditions by subject-specific decoders (Panel A) and group decoders (Panel B). Trained and Test data were from matching listening conditions.

## Discussion

### Intelligibility Boosts Speech Envelope Reconstruction Accuracy, But the Effect is Small

A central question in speech perception research is what neural tracking of the amplitude envelope can disclose about comprehension (Ding & Simon, 2014). For the purposes of clinically assessing auditory speech processing, an equally pertinent question is whether comprehension contributes to neural tracking (our first hypothesis). This is relevant where auditory input is likely to be severely spectrally degraded, as would be the case for CI patients. For example, an individual with relatively poor spectral resolution might nonetheless show good neural speech tracking under optimal listening conditions. Speech intelligibility could, therefore, obscure real problems occurring early in the auditory processing hierarchy, such as issues at the electrode-neural interface, or sub-optimal device configuration. Conversely, if accurate decoding is dependent on speech comprehension, this would prevent its use as an objective measure of auditory encoding in many clinical populations. Hence, it is crucial to understand the relationship between speech stimulus clarity and intelligibility as one of many endogenously originating factors that could shape early sensory processing. In the current study, we address this challenge by presenting participants with intelligible and non-intelligible speech at varying levels of spectral degradation. We attempted to minimise the influence of stimulus properties that were previously confounded with intelligibility, including acoustic properties, speaker and voice characteristics, and auditory attention. We, moreover, quantify the generalisability of speech decoding in the context of our experimental manipulations, as well as across subject-specific versus group-level reconstruction techniques. Our findings cohere with an emerging consensus that envelope tracking is possible without speech understanding (Baltzell et al., 2017; Gillis et al., 2023; Karunathilake et al., 2023; Kösem et al., 2023), yet intelligibility nonetheless exerts an enhancing effect on decoding accuracy in the current dataset and thereby supports our first hypothesis.

Our linear mixed modelling results suggest the effect of intelligibility is consistent, but small. Importantly, we found no interaction between spectral degradation and intelligibility on speech tracking and thereby no support for our second hypothesis, although key questions remain. One relates to the methodology itself, namely, the issue of decoder significance. Although our analysis and hypotheses focused on decoding accuracy across participants, a clinician may be more interested in decoder performance at the patient level. Our descriptive analysis showed that the rank of the observed decoder accuracy relative to its null distribution was inconsistent across decoding methods and listening conditions. Future work with inferential statistics should confirm our observations here, but it is potentially problematic that only English unprocessed speech was decoded significantly for all participants. Given that the permuted feature, the amplitude envelope, is largely unchanged by Spectral Degradation and appears to share basic temporal properties across English and Dutch, fluctuations in the false positive rate (i.e., where a randomly permuted envelope is decoded with higher accuracy than the true stimulus envelope) are presumably attributable to differences in the listener’s neural response. It is possible that some brain states are more amenable to spurious correlations with the speech envelope: We can envision this possibly occurring due to enhanced oscillatory activity at the syllable rate, as one hypothetical example.

Another unresolved question is whether individual measures of comprehension can be predicted by speech envelope decoding. We observed a tenuous link between self-reported comprehension and subject-specific neural decoding, but its magnitude was reduced when accounting for the brain response to spectral degradation, independent of understanding. One limitation in our study is that we did not collect any objective measure of speech comprehension, such as asking participants about the narrative content of the story. This decision was made partly in the interest of time, and partly because an equivalent measure was not available for Dutch listening conditions. It is possible that participants’ self-described ability to follow is less informative than comprehension questions or word reports. Another consideration is that, although English and Dutch are mutually unintelligible, they do share many suprasegmental properties in common. As we wished to minimise acoustic and linguistic differences between intelligible and non-intelligible conditions, this was a deliberate choice for our experimental design. Prosody, although not necessarily lexically informative, nonetheless conveys salient information, ranging from the rhythm and structure of words and phrases, to emotional affect, to the pacing of action and dialogue. Hence, although listeners in our study reported very little understanding or sense of engagement relating to the Dutch stimuli, our decoding results probably still capture language- and speech-specific neural responses beyond strictly acoustic processing (Myers et al., 2019).

### Equivalent Envelope Reconstruction of Unprocessed and Spectrally Degraded Speech

Based on the literature, we formed different hypotheses for spectral degradation in the current study. In one scenario, the neural decoding of Dutch speech worsens with declining acoustic clarity, possibly because listeners are unable to benefit from semantic prediction or other top-down processes. As it turned out, at the group level, speech tracking level was similar across unprocessed, vocoded, and vocoded speech with additional blurring distortion. Given that the broadband amplitude envelope was more or less identical across spectral degradation conditions, our result speaks to the dominance of this acoustic feature in the speech-evoked EEG (Ding & Simon, 2014). For example, Prinsloo and Lalor (2022) showed that, even for artificial speech stimuli where the envelope conveys no meaningful information, it evokes a stronger cortical response in listeners than competing acoustic features that do facilitate comprehension. One aspect in which our study differs from others is that we did not manipulate spectral resolution to the point where speech could no longer be understood at all. This preserves the orthogonality between acoustic quality and speech understanding, but challenges straightforward comparison to previous results. For instance, whereas some authors reported no differences in the neural response to unprocessed and 4-channel vocoded speech (Dong & Gai, 2021), other groups did see decoding accuracy decrease when speech was vocoded with five or fewer channels (Chen et al., 2023). Hence, speech may not have been spectrally degraded enough in the current experiment to elicit differences in decoding accuracy, assuming it is indeed spectral resolution and not intelligibility that impacted stimulus reconstruction (Chen et al., 2023). It is, however, possible that we would observe spatial and temporal differences when characterising the EEG response itself. This idea is supported by evidence from studies that used encoders and other models wherein weights and time lags are representative of neural activation patterns, and can, therefore, be interpreted by researchers (Holdgraf et al., 2017), noting that these additional parameters–some of which are manually selected– could complicate their use in clinical settings. For example, behaviourally measured comprehension was the same for unprocessed and 16-channel vocoded speech, but the modelled neural response was more similar between 16-channel and 8-channel vocoded speech than between unprocessed and 16-channel vocoded speech (Kong et al., 2015). Another study employing invasive stereoelectroencephalography (sEEG) found an early high-gamma (60 − 140 Hz) power response, localised to primary auditory cortex, that responded preferentially to vocoded speech, and a later theta (5 − 8 Hz) response, localised to nonprimary auditory areas mainly in the right hemisphere, that responded preferentially to unprocessed speech (Xu et al., 2023). In addition to characterising the temporal profile or waveform of the neural response, there are other measures that reveal differences between unprocessed and vocoded speech processing, such as speech-brain coherence, which describes the phase relationship between stimulus features and the neural response. Studies employing this technique suggest that vocoding speech shifts the coherence centre frequency from the syllabic rate (∼4 Hz) towards the slightly faster acoustic modulation rate (∼5 Hz; Schmidt et al., 2021; Chen et al., 2023).Together, these findings imply the possibility of distinct auditory processing pathways for unprocessed and vocoded speech, despite their equivalent amenability to neural decoding. In particular, Xu and colleagues (2023) argued that the sEEG data reveal a functional dissociation between automatic processing of low-level acoustic information, and a slightly later response that is selective for intelligible speech; however, as speech intelligibility was fully confounded with acoustic clarity in their study, it is unknown whether spatial-temporal changes in the neural response are the result of waning comprehension or increasing spectral degradation. A shift in perceptual attention from one putative system to the other could, in theory, contribute to the sudden “pop-out” listeners sometimes experience when hearing vocoded speech (Corcoran et al., 2023; Davis et al., 2005; Karunathilake et al., 2023; Kösem et al., 2023). Presumably, under ecologically relevant conditions, both the automatic acoustic and speech-selective systems would continuously interact at multiple stages of auditory processing. Of relevance to our results, it is possible that the early, acoustically driven response contributed to statistical significance detection in subject-specific decoding of Dutch speech, which tended to improve with decreasing spectral resolution.

### Decoding Within and Across Individuals and Listening Conditions

By training and testing within and across listening conditions, we were able to quantify the generalisability of speech tracking. Although subject-specific and group-level decoding showed good within-individual correlation, group decoders produced comparably low accuracy values. They were also less discriminating with respect to listening condition, to the extent that there was a minimal effect of training the group decoders on English versus Dutch speech. Approximately one third of group decoders, furthermore, failed to meet statistical significance according to random permutation testing. When comparing decoding across the entire experiment (Figure 5), it can be seen that some individuals particularly benefit from subject-specific decoding, at least with regards to accuracy. Potential clinical implications, however, will also depend on the relative robustness of decoding–it is possible that subject-specific decoders are overfit–and the ability to capture specific aspects of the neural response that are relevant to speech outcomes. While these preliminary results do not support the clinical application of neural decoding to assess fine spectral detail or estimate speech comprehension, both subject- and group-level decoders showed reliable tracking for the majority of participants, and could therefore be used as a more fundamental measure of auditory speech processing.

As the acoustic quality of test speech becomes less similar to training speech, decoding accuracy wanes. This pattern is symmetrical across the three levels of spectral degradation, with no apparent advantage unique to training and testing on unprocessed speech. That result stands in contrast to intelligibility, where we do record consistently better decoding for the language that is understood, regardless of whether the model was trained on intelligible speech. There was, however, substantial variability in the neural response to spectral degradation between participants (Gransier et al., 2021). This is evidenced in the current data set by the wide confidence intervals for the model estimates and effect sizes of spectral degradation-related predictors. Given the individuality and diversity of neural responses observed in the present dataset, which was contributed by typically hearing younger adults, it seems unlikely that a single normative or “healthy” decoder would be suitable for clinical or non-neurotypical populations–at least for the purposes of assessing sensory-acoustic clarity in speech processing.

Our findings do underscore the potential clinical viability of subject-specific decoding, for which statistically significant decoders can be trained using ∼10 minutes of data per listening condition. Although the absolute magnitude of neural decoding accuracy is unlikely to be meaningful at the population level, we can envision a paradigm wherein speech or speech-like stimuli are manipulated on a specific acoustic dimension. Establishing the patient’s individual threshold at which this manipulation can be detected in the EEG response could form the basis of an objective measure of auditory processing, for example (Goehring et al., 2020, 2021). Given the observed substantial variability across spectral degradation conditions within individuals, it seems unlikely that neural decoding can reveal differences in spectral resolution for individual CI listeners when predicting the speech amplitude envelope. Still, it may serve as a principal measure of speech envelope perception during naturalistic speech listening independent of speech understanding or spectral resolution and therefore be helpful in the CI device fitting process. CI speech processing relies fully on the successful transmission of speech envelope information across a limited number of frequency channels and an objective measure of this capacity could serve as a crucial tool to support the device fitting and validation process in clinics. It may be particularly useful for patients with limited verbal communication skills, for whom device fitting can be very challenging. The previously mentioned CI stimulation artefacts complicate EEG-based measurements in CI listeners, but methods to mitigate CI-related artefacts with the use of stimulation gaps (Somers et al., 2018), computational artefact removal (Intartaglia et al., 2022) or interleaved stimulation (Carlyon et al., 2021, Guerit et al., 2023) have recently been proposed and successfully applied to measure neural responses in CI listeners with EEG. A combination of artefact removal techniques with objective, EEG-based measures of speech transmission warrant further research but represent a promising venue to develop future clinical applications.

### Conclusion

An objective measure of neural speech encoding function would benefit patients and clinicians, as well as researchers involved in hearing sciences or the cognitive neuroscience of speech. However, for an objective measure to be useful, it should also work for individuals for whom the speech signal is sparse or distorted, such as CI listeners. Thus far, efforts to link physiology with subjective and objective behavioural measures of speech perception have been limited by confounding factors, such as acoustic clarity and prior knowledge. In this study, we have attempted to control these and other dimensions that may covary with intelligibility. The results show that, at the sample level, the amplitude envelope is robustly represented within the neural response to speech, even when speech is not understood, and that this response can be detected in the context of severe spectral degradation. However, we also find limited evidence for a relationship between speech comprehension and neural decoding of the envelope, with slightly but significantly higher values in the intelligible language. We also observe substantial between-participant variation in the brain response to different listening conditions. It is possible that within-participant analyses, whether by permutation testing, or presenting listeners with parameterised stimuli, may yield greater insights into hearing health and speech reception than the interpretation of the absolute accuracy of neural decoding *per se*. We conclude that while more research is warranted to understand the sources of variability observed in these data, neural decoding of the speech envelope may serve as a basic objective measure of auditory speech processing during CI listening and should be further investigated.

## Acknowledgments

This research was funded by the Wellcome Trust Institutional Strategic Support Fund (ADM) and the Medical Research Council UK Career Development Award Fellowship MR/T03095X/1 (TG). For the purpose of open access, the authors have applied a Creative Commons Attribution (CC BY) license to any Author Accepted Manuscript version arising.

## Appendix

### Reaction Times

**Table A1.** Model selection for the dependent variable of Median Reaction Times (s). Missing values of AIC indicate the model did not converge.

| Model | df | AIC | Formula |
| --- | --- | --- | --- |
| 1 | 3 | -9.89 | 1 + (1 \| Participant) |
| 2 | 5 | -45 | 1 + Spectral Degradation + (1 \| Participant) |
| 3 | 4 | -46.05 | 1 + Language + (1 \| Participant) |
| 4 | 4 | -60.82 | 1 + Ability to Follow + (1 \| Participant) |
| 5 | 4 | -57.57 | 1 + Engagement + (1 \| Participant) |
| 6 | 6 | -95.71 | 1 + Spectral Degradation + Language + (1 \| Participant) |
| 7 | 6 | -91.46 | 1 + Spectral Degradation + Ability to Follow + (1 \| Participant) |
| 8 | 8 | -83.51 | 1 + Spectral Degradation * Language + (1 \| Participant) |
| 9 | 11 | — | 1 + Spectral Degradation + Language + (1 + Spectral Degradation \| Participant) |
| 10 | 8 | -99.83 | 1 + Spectral Degradation + Language + (1 + Language \| Participant) |
| 11 | 14 | — | 1 + Spectral Degradation + Language + (1 + Spectral Degradation \| Participant) + (Language \| Participant) |

**Table A2.** Model formula: Median RT ∼ 1 + Spectral Degradation + Language + (1 + Language | Participant). Standardized parameters were obtained by fitting the model on a standardized version of the dataset. 95% Confidence Intervals (CIs) and p-values were computed using a Wald t-distribution with Kenward-Roger approximation.

| Parameter | Estimate | 95% CI | *t* | df | *p* | Std. Estimate | Std. Estimate 95% CI |
| --- | --- | --- | --- | --- | --- | --- | --- |
| Intercept | 1.02 | [0.94, 1.10] | 25.33 | 45.28 | < 0.001 | -0.56 | [-0.84, -0.28] |
| Vocoded | 0.05 | [0.01, 0.10] | 2.43 | 150.00 | 0.016 | 0.18 | [0.03, 0.33] |
| Vocoded + Blurring | 0.20 | [0.16, 0.24] | 9.21 | 150.00 | < 0.001 | 0.69 | [0.54, 0.83] |
| Dutch | 0.16 | [0.11, 0.21] | 6.50 | 37.00 | < 0.001 | 0.55 | [0.38, 0.72] |
| **Random Effects** |  |  |  |  |  |  |  |
| Within-Group Variance (SD) |  | 0.02 (0.13) |  |  |  |  |  |
| Between-Group Variance (SD) |  |  |  |  |  |  |  |
|  | Intercept (Participant) | 0.05 (0.22) |  |  |  |  |  |
|  | Slope (Language) | 0.01 (0.11) |  |  |  |  |  |
| Correlations |  |  |  |  |  |  |  |
|  | Participant × Language | 0.01 |  |  |  |  |  |
| Groups |  |  |  |  |  |  |  |
|  | Participant | 38 |  |  |  |  |  |
| Observations |  | 228 |  |  |  |  |  |
| **Model Metrics** |  |  |  |  |  |  |  |
| AIC | BIC | R<sup>2</sup> (Conditional) | R<sup>2</sup> (Marginal) | ICC | RMSE |  |  |
| -99.83 | -72.40 | 0.79 | 0.16 | 0.75 | 0.12 |  |  |

**Table A3.**
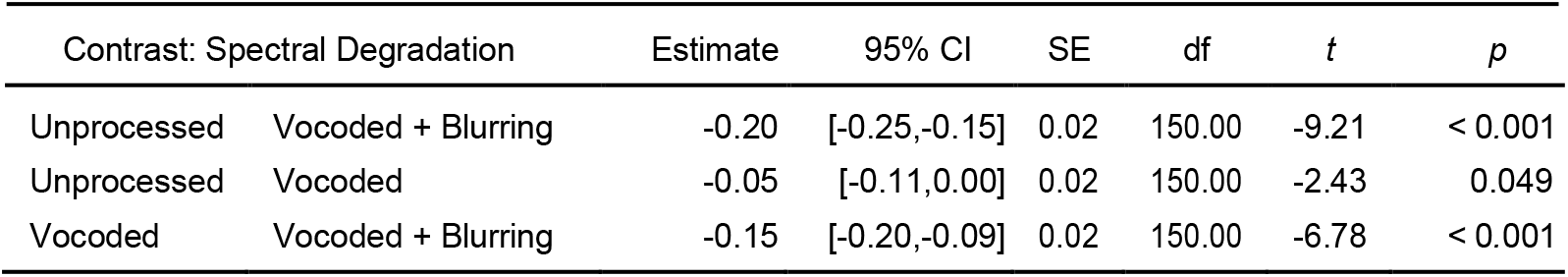
Contrast of Median Reaction Times (s) estimated marginal means for Spectral Degradation. P-values are adjusted using the Bonferroni method.

| Contrast: Spectral Degradation |  | Estimate | 95% CI | SE | df | *t* | *p* |
| --- | --- | --- | --- | --- | --- | --- | --- |
| Unprocessed | Vocoded + Blurring | -0.20 | [-0.25, -0.15] | 0.02 | 150.00 | -9.21 | < 0.001 |
| Unprocessed | Vocoded | -0.05 | [-0.11, 0.00] | 0.02 | 150.00 | -2.43 | 0.049 |
| Vocoded | Vocoded + Blurring | -0.15 | [-0.20, -0.09] | 0.02 | 150.00 | -6.78 | < 0.001 |

**Table A4.** Model selection for the dependent variable of subject-specific Decoding Accuracy (r). Missing values of AIC indicate the model did not converge.

| Model | df | AIC | Formula |
| --- | --- | --- | --- |
| 1 | 3 | -3898.05 | 1 + (1 \| Participant) |
| 2 | 5 | -3885.63 | 1 + Trained: Spectral Degradation + (1 \| Participant) |
| 3 | 5 | -3913.99 | 1 + Test: Spectral Degradation + (1 \| Participant) |
| 4 | 7 | -3901.82 | 1 + Trained: Spectral Degradation + Test: Spectral Degradation + (1 \| Participant) |
| 5 | 11 | -3962.91 | 1 + Trained: Spectral Degradation × Test: Spectral Degradation + (1 \| Participant) |
| 6 | 16 | — | 1 + Trained: Spectral Degradation × Test: Spectral Degradation + (Trained: Spectral Degradation \| Participant) |
| 7 | 16 | -4326.58 | 1 + Trained: Spectral Degradation × Test: Spectral Degradation + (Test: Spectral Degradation \| Participant) |
| 8 | 17 | -4317.53 | 1 + Trained: Spectral Degradation × Test: Spectral Degradation + Trained: Language + (Test: Spectral Degradation \| Participant) |
| 9 | 17 | -4367.24 | 1 + Trained: Spectral Degradation × Test: Spectral Degradation + Test: Language + (Test: Spectral Degradation \| Participant) |
| 10 | 18 | -4358.28 | 1 + Trained: Spectral Degradation × Test: Spectral Degradation + Trained: Language + Test: Language + (Test: Spectral Degradation \| Participant) |
| 11 | 19 | -4373.39 | 1 + Trained: Spectral Degradation × Test: Spectral Degradation + Trained: Language × Test: Language + (Test: Spectral Degradation \| Participant) |
| 12 | 22 | — | 1 + Trained: Spectral Degradation × Test: Spectral Degradation + Trained: Language × Test: Language + (Test: Spectral Degradation \| Participant) + (Trained: Language \| Participant) |
| 13 | 22 | — | 1 + Trained: Spectral Degradation × Test: Spectral Degradation + Trained: Language × Test: Language + (Test: Spectral Degradation \| Participant) + (Test: Language \| Participant) |
| 14 | 27 | -4310.04 | 1 + Trained: Spectral Degradation × Test: Spectral Degradation × Trained: Language + Trained: Language × Test: Language + (Test: Spectral Degradation \| Participant) |
| 15 | 27 | -4319.79 | 1 + Trained: Spectral Degradation × Test: Spectral Degradation × Test: Language + Trained: Language × Test: Language + (Test: Spectral Degradation \| Participant) |
| 16 | 43 | -4192.12 | 1 + Trained: Spectral Degradation × Test: Spectral Degradation × Trained: Language × Test: Language + (Test: Spectral Degradation \| Participant) |

### Subject-Specific Decoding

**Table A5.**
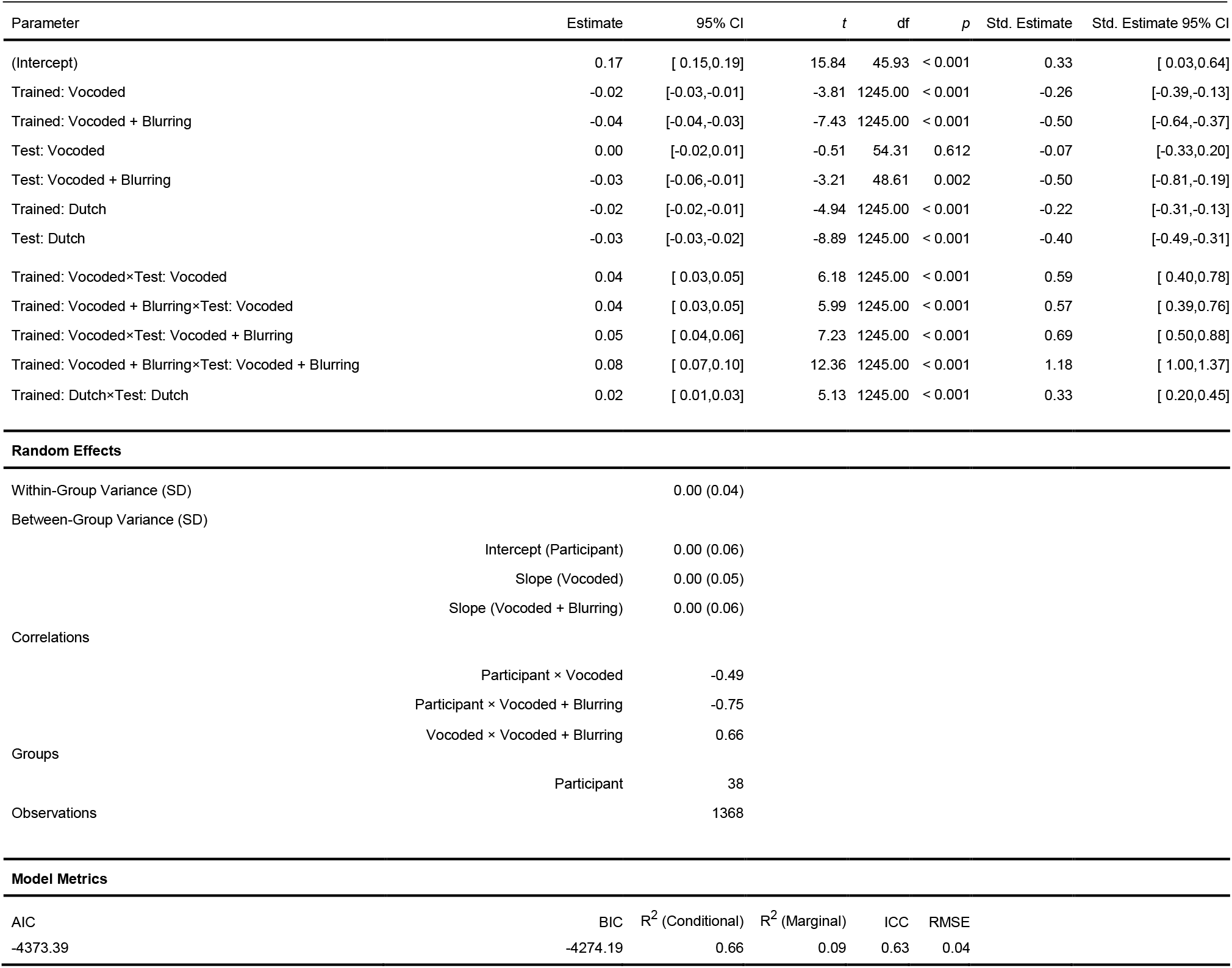
Model formula: Decoding Accuracy R Value ∼ 1 + Trained: Spectral Degradation × Test: Spectral Degradation + Trained: Language × Test: Language + (Test: Spectral Degradation | Participant). Standardized parameters were obtained by fitting the model on a standardized version of the dataset. 95% Confidence Intervals (CIs) and p-values were

**Table A6.** Contrast of subject-specific Decoding Accuracy (r) estimated marginal means for Test: Spectral Degradation within levels of Trained: Spectral Degradation. P-values are adjusted using the Bonferroni method.

| Contrast: Test: Spectral Degradation |  | At: Trained: Spectral Degradation | Estimate | 95% CI | SE | df | t | p |
| --- | --- | --- | --- | --- | --- | --- | --- | --- |
| Unprocessed | Vocoded | Unprocessed | 0.00 | [-0.02,0.03] | 0.01 | 54.31 | 0.51 | 1.000 |
| Unprocessed | Vocoded + Blurring | Unprocessed | 0.03 | [ 0.01,0.06] | 0.01 | 48.61 | 3.21 | 0.007 |
| Vocoded | Vocoded + Blurring | Unprocessed | 0.03 | [ 0.01,0.05] | 0.01 | 56.33 | 3.39 | 0.004 |
| Unprocessed | Vocoded | Vocoded | -0.04 | [-0.06,-0.01] | 0.01 | 54.31 | -3.97 | 0.001 |
| Unprocessed | Vocoded + Blurring | Vocoded | -0.01 | [-0.04,0.01] | 0.01 | 48.61 | -1.26 | 0.637 |
| Vocoded | Vocoded + Blurring | Vocoded | 0.02 | [ 0.00,0.05] | 0.01 | 56.33 | 2.59 | 0.036 |
| Unprocessed | Vocoded | Vocoded + Blurring | -0.04 | [-0.06,-0.01] | 0.01 | 54.31 | -3.83 | 0.001 |
| Unprocessed | Vocoded + Blurring | Vocoded + Blurring | -0.05 | [-0.08,-0.02] | 0.01 | 48.61 | -4.45 | < 0.001 |
| Vocoded | Vocoded + Blurring | Vocoded + Blurring | -0.01 | [-0.03,0.01] | 0.01 | 56.33 | -1.43 | 0.478 |

**Table A7.** Contrast of subject-specific Decoding Accuracy (r) estimated marginal means for Test: Language within levels of Trained: Language. P-values are adjusted using the Bonferroni method.

| Contrast: Test: Language |  | At: Trained: Language | Estimate | 95% CI | SE | df | t | p |
| --- | --- | --- | --- | --- | --- | --- | --- | --- |
| English | Dutch | English | 0.03 | [0.02,0.03] | 0.00 | 1245.00 | 8.89 | < 0.001 |
| English | Dutch | Dutch | 0.01 | [0.00,0.01] | 0.00 | 1245.00 | 1.63 | 0.104 |

### Group Decoding

**Table A8.** Model selection for the dependent variable of group Decoding Accuracy (r). Missing values of AIC indicate the model did not converge.

| Model | df | AIC | Formula |
| --- | --- | --- | --- |
| 1 | 3 | -4050.93 | 1 + (1 \| Participant) |
| 2 | 5 | -4035.82 | 1 + Trained: Spectral Degradation + (1 \| Participant) |
| 3 | 5 | -4053.48 | 1 + Test: Spectral Degradation + (1 \| Participant) |
| 4 | 7 | -4038.48 | 1 + Trained: Spectral Degradation + Test: Spectral Degradation + (1 \| Participant) |
| 5 | 11 | -4040.59 | 1 + Trained: Spectral Degradation × Test: Spectral Degradation + (1 \| Participant) |
| 6 | 16 | — | 1 + Trained: Spectral Degradation × Test: Spectral Degradation + (Trained: Spectral Degradation \| Participant) |
| 7 | 16 | -4473.73 | 1 + Trained: Spectral Degradation × Test: Spectral Degradation + (Test: Spectral Degradation \| Participant) |
| 8 | 17 | -4462.27 | 1 + Trained: Spectral Degradation × Test: Spectral Degradation + Trained: Language + (Test: Spectral Degradation \| Participant) |
| 9 | 17 | -4509.57 | 1 + Trained: Spectral Degradation × Test: Spectral Degradation + Test: Language + (Test: Spectral Degradation \| Participant) |
| 10 | 20 | — | 1 + Trained: Spectral Degradation × Test: Spectral Degradation + Test: Language + (Test: Spectral Degradation \| Participant) + (Test: Language \| Participant) |
| 11 | 19 | -4495.02 | 1 + Trained: Spectral Degradation × Test: Spectral Degradation + Trained: Language × Test: Language + (Test: Spectral Degradation \| Participant) |
| 12 | 43 | -4295.10 | 1 + Trained: Spectral Degradation × Test: Spectral Degradation × Trained: Language × Test: Language + (Test: Spectral Degradation \| Participant) |

**Table A9.** Model formula: Decoding Accuracy R Value ∼ 1 + Trained: Spectral Degradation × Test: Spectral Degradation + Test: Language + (Test: Spectral Degradation | Participant). Standardized parameters were obtained by fitting the model on a standardized version of the dataset. 95% Confidence Intervals (CIs) and p-values were computed using a Wald t-distribution with Kenward-Roger approximation.

| Parameter | Estimate | 95% CI | *t* | df | *p* | Std. Estimate | Std. Estimate 95% CI |
| --- | --- | --- | --- | --- | --- | --- | --- |
| (Intercept) | 0.11 | [ 0.10,0.13] | 16.28 | 53.39 | < 0.001 | 0.22 | [-0.01,0.44] |
| Trained: Vocoded | -0.01 | [-0.02,0.00] | -2.26 | 1247.00 | 0.024 | -0.17 | [-0.31,-0.02] |
| Trained: Vocoded + Blurring | -0.03 | [-0.04,-0.02] | -5.85 | 1247.00 | < 0.001 | -0.43 | [-0.57,-0.29] |
| Test: Vocoded | 0.00 | [-0.02,0.02] | -0.28 | 50.28 | 0.778 | -0.04 | [-0.36,0.27] |
| Test: Vocoded + Blurring | -0.03 | [-0.05,0.00] | -2.46 | 48.08 | 0.018 | -0.42 | [-0.76,-0.08] |
| Test: Dutch | -0.02 | [-0.02,-0.01] | -7.01 | 1247.00 | < 0.001 | -0.24 | [-0.31,-0.17] |
| Trained: Vocoded×Test: Vocoded | 0.03 | [ 0.01,0.04] | 3.89 | 1247.00 | < 0.001 | 0.40 | [ 0.20,0.61] |
| Trained: Vocoded + Blurring×Test: Vocoded | 0.03 | [ 0.02,0.05] | 5.04 | 1247.00 | < 0.001 | 0.52 | [ 0.32,0.73] |
| Trained: Vocoded×Test: Vocoded + Blurring | 0.03 | [ 0.02,0.05] | 5.23 | 1247.00 | < 0.001 | 0.54 | [ 0.34,0.75] |
| Trained: Vocoded + Blurring×Test: Vocoded + Blurring | 0.05 | [ 0.04,0.07] | 8.17 | 1247.00 | < 0.001 | 0.85 | [ 0.64,1.05] |
| **Random Effects** |  |  |  |  |  |  |  |
| Within-Group Variance (SD) |  | 0.00 (0.04) |  |  |  |  |  |
| Between-Group Variance (SD) |  |  |  |  |  |  |  |
|  | Intercept (Participant) | 0.00 (0.04) |  |  |  |  |  |
|  | Slope (Vocoded) | 0.00 (0.05) |  |  |  |  |  |
|  | Slope (Vocoded + Blurring) | 0.00 (0.06) |  |  |  |  |  |
| Correlations |  |  |  |  |  |  |  |
|  | Participant × Vocoded | -0.54 |  |  |  |  |  |
|  | Participant × Vocoded + Blurring | -0.45 |  |  |  |  |  |
|  | Vocoded × Vocoded + Blurring | 0.64 |  |  |  |  |  |
| Groups |  |  |  |  |  |  |  |
|  | Participant | 38 |  |  |  |  |  |
| Observations |  | 1368 |  |  |  |  |  |
| **Model Metrics** |  |  |  |  |  |  |  |
| AIC | BIC | R<sup>2</sup> (Conditional) | R<sup>2</sup> (Marginal) | ICC | RMSE |  |  |
| -4509.57 | -4420.82 | 0.60 | 0.05 | 0.57 | 0.04 |  |  |

**Table A10.** Contrast of group Decoding Accuracy (r) estimated marginal means for Test: Spectral Degradation within levels of Trained: Spectral Degradation. P-values are adjusted using the Bonferroni method.

| Contrast: Test: Spectral Degradation |  | At: Trained: Spectral Degradation | Estimate | 95% CI | SE | df | t | p |
| --- | --- | --- | --- | --- | --- | --- | --- | --- |
| Unprocessed | Vocoded | Unprocessed | 0.00 | [-0.02,0.03] | 0.01 | 50.28 | 0.28 | 1.000 |
| Unprocessed | Vocoded + Blurring | Unprocessed | 0.03 | [ 0.00,0.05] | 0.01 | 48.08 | 2.46 | 0.053 |
| Vocoded | Vocoded + Blurring | Unprocessed | 0.02 | [ 0.00,0.05] | 0.01 | 53.41 | 2.57 | 0.039 |
| Unprocessed | Vocoded | Vocoded | -0.02 | [-0.05,0.00] | 0.01 | 50.28 | -2.26 | 0.085 |
| Unprocessed | Vocoded + Blurring | Vocoded | -0.01 | [-0.03,0.02] | 0.01 | 48.08 | -0.72 | 1.000 |
| Vocoded | Vocoded + Blurring | Vocoded | 0.01 | [-0.01,0.04] | 0.01 | 53.41 | 1.61 | 0.337 |
| Unprocessed | Vocoded | Vocoded + Blurring | -0.03 | [-0.05,-0.01] | 0.01 | 50.28 | -3.01 | 0.012 |
| Unprocessed | Vocoded + Blurring | Vocoded + Blurring | -0.03 | [-0.05,0.00] | 0.01 | 48.08 | -2.50 | 0.047 |
| Vocoded | Vocoded + Blurring | Vocoded + Blurring | 0.00 | [-0.02,0.03] | 0.01 | 53.41 | 0.34 | 1.000 |

## References

Aiken, S. J., & Picton, T. W. (2008). Human cortical responses to the speech envelope. Ear and Hearing, 29(2), 139–157.

Alikovic, E., Ng, E. H. N., Fiedler, L., Santurette, S., Innes-Brown, H., & Graversen, C. (2021). Effects of hearing aid noise reduction on early and late cortical representations of competing talkers in noise. Frontiers in Neuroscience, 15, 636060.

Aljarboa, G. S., Bell, S. L., & Simpson, D. M. (2023). Detecting cortical responses to continuous running speech using eeg data from only one channel. International Journal of Audiology, 62(3), 199–208.

Asilador, A., & Llano, D. A. (2021). Top-down inference in the auditory system: potential roles for corticofugal projections. Frontiers in Neural Circuits, 14, 615259.

Baltzell, L. S., Horton, C., Shen, Y., Richards, V. M., D’Zmura, M., & Srinivasan, R. (2016). Attention selectively modulates cortical entrainment in different regions of the speech spectrum. Brain Research, 1644, 203–212.

Baltzell, L. S., Srinivasan, R., & Richards, V. M. (2017). The effect of prior knowledge and intelligibility on the cortical entrainment response to speech. Journal of Neurophysiology, 118(6), 3144–3151.

Barajas, M. C. O., Guevara, R., & Gervain, J. (2021). The origins and development of speech envelope tracking during the first months of life. Developmental Cognitive Neuroscience, 48, 100915.

Bates, D., Mächler, M., Bolker, B., & Walker, S. (2014). Fitting linear mixed-effects models using lme4. arXiv preprint arXiv:1406.5823.

Biesmans, W., Das, N., Francart, T., & Bertrand, A. (2016). Auditory-inspired speech envelope extraction methods for improved eeg-based auditory attention detection in a cocktail party scenario. IEEE Transactions on Neural Systems and Rehabilitation Engineering, 25(5), 402–412.

Bingabr, M., Espinoza-Varas, B., & Loizou, P. C. (2008). Simulating the effect of spread of excitation in cochlear implants. Hearing Research, 241(1-2), 73–79.

Boisvert, I., Reis, M., Au, A., Cowan, R., & Dowell, R. C. (2020). Cochlear implantation outcomes in adults: A scoping review. PLoS One, 15(5), e0232421.

Brodbeck, C., & Simon, J. Z. (2020). Continuous speech processing. Current Opinion in Physiology, 18, 25–31.

Broderick, M. P., Anderson, A. J., Di Liberto, G. M., Crosse, M. J., & Lalor, E. C. (2018). Electrophysiological correlates of semantic dissimilarity reflect the comprehension of natural, narrative speech. Current Biology, 28(5), 803–809.

Burle, B., Spieser, L., Roger, C., Casini, L., Hasbroucq, T., & Vidal, F. (2015). Spatial and temporal resolutions of eeg: Is it really black and white? a scalp current density view. International Journal of Psychophysiology, 97(3), 210–220.

Burnham, K. P., Anderson, D. R., & Huyvaert, K. P. (2011). Aic model selection and multimodel inference in behavioral ecology: Some background, observations, and comparisons. Behavioral Ecology and Sociobiology, 65, 23–35.

Byrne, D., Dillon, H., Tran, K., Arlinger, S., Wilbraham, K., Cox, R., … & Ludvigsen, C. (1994). An international comparison of long-term average speech spectra. The Journal of the Acoustical Society of America, 96(4), 2108–2120.

Carlyon, R. P., & Goehring, T. (2021). Cochlear implant research and development in the twenty-first century: A critical update. Journal of the Association for Research in Otolaryngology, 22(5), 481–508.

Carlyon, R. P., Guérit, F., Deeks, J. M., Harland, A., Gransier, R., Wouters, J., … & Bance, M. (2021). Using interleaved stimulation to measure the size and selectivity of the sustained phase-locked neural response to cochlear implant stimulation. Journal of the Association for Research in Otolaryngology, 22, 141–159.

Casas, A. S. H., Lajnef, T., Pascarella, A., Guiraud-Vinatea, H., Laaksonen, H., Bayle, D., Jerbi, K., & Boulenger, V. (2021). Neural oscillations track natural but not artificial fast speech: Novel insights from speech-brain coupling using meg. NeuroImage, 244, 118577.

Chen, Y.-P., Schmidt, F., Keitel, A., Rösch, S., Hauswald, A., & Weisz, N. (2023). Speech intelligibility changes the temporal evolution of neural speech tracking. NeuroImage, 268, 119894.

Christophe, A., & Morton, J. (1998). Is dutch native english? linguistic analysis by 2-month-olds. Developmental Science, 1(2), 215–219.

Cogan, G. B., & Poeppel, D. (2011). A mutual information analysis of neural coding of speech by low-frequency MEG phase information. Journal of Neurophysiology, 106(2), 554–563.

Corcoran, A. W., Perera, R., Koroma, M., Kouider, S., Hohwy, J., & Andrillon, T. (2023). Expectations boost the reconstruction of auditory features from electrophysiological responses to noisy speech. Cerebral Cortex, 33(3), 691– 708.

Crosse, M. J., Butler, J. S., & Lalor, E. C. (2015). Congruent visual speech enhances cortical entrainment to continuous auditory speech in noise-free conditions. Journal of Neuroscience, 35(42), 14195–14204.

Crosse, M. J., Di Liberto, G. M., Bednar, A., & Lalor, E. C. (2016). The multivariate temporal response function (mtrf) toolbox: A matlab toolbox for relating neural signals to continuous stimuli. Frontiers in Human Neuroscience, 10, 604.

Crosse, M. J., Zuk, N. J., Di Liberto, G. M., Nidiffer, A. R., Molholm, S., & Lalor, E. C. (2021). Linear modeling of neurophysiological responses to speech and other continuous stimuli: Methodological considerations for applied research. Frontiers in Neuroscience, 1350.

Cychosz, M., Winn, M., & Goupell, M. J. (2023). How (not) to vocode: Using channel vocoders for cochlear-implant research. 10.31234/osf.io/yrqnu

Davis, M. H., & Johnsrude, I. S. (2007). Hearing speech sounds: Top-down influences on the interface between audition and speech perception. Hearing Research, 229(1-2), 132–147.

Davis, M. H., Johnsrude, I. S., Hervais-Adelman, A., Taylor, K., & McGettigan, C. (2005). Lexical information drives perceptual learning of distorted speech: Evidence from the comprehension of noise-vocoded sentences. Journal of Experimental Psychology: General, 134(2), 222.

De Clercq, P., Vanthornhout, J., Vandermosten, M., & Francart, T. (2023). Beyond linear neural envelope tracking: A mutual information approach. Journal of Neural Engineering, 20(2), 026007.

de Cheveigné, A., & Arzounian, D. (2018). Robust detrending, rereferencing, outlier detection, and inpainting for multichannel data. NeuroImage, 172, 903–912.

Decruy, L., Vanthornhout, J., & Francart, T. (2020). Hearing impairment is associated with enhanced neural tracking of the speech envelope. Hearing Research, 393, 107961.

Di Liberto, G. M., O’Sullivan, J. A., & Lalor, E. C. (2015). Low-frequency cortical entrainment to speech reflects phoneme-level processing. Current Biology, 25(19), 2457–2465.

Ding, N., Chatterjee, M., & Simon, J. Z. (2014). Robust cortical entrainment to the speech envelope relies on the spectro-temporal fine structure. NeuroImage, 88, 41–46.

Ding, N., & Simon, J. Z. (2012). Emergence of neural encoding of auditory objects while listening to competing speakers. Proceedings of the National Academy of Sciences, 109(29), 11854–11859.

Ding, N., & Simon, J. Z. (2014). Cortical entrainment to continuous speech: Functional roles and interpretations. Frontiers in Human Neuroscience, 8, 311.

Dolhopiatenko, H., & Nogueira, W. (2023). Selective attention decoding in bimodal cochlear implant users. Frontiers in Neuroscience, 16, 1057605.

Dong, Y., & Gai, Y. (2021). Speech perception with noise vocoding and background noise: An eeg and behavioral study. Journal of the Association for Research in Otolaryngology, 22, 349–363.

Doyle, A. C. (1903). De terugkeer van sherlock holmes [Available at https://www.gutenberg.org/ebooks/29490. Accessed on September 08, 2023]. https://www.gutenberg.org/ebooks/29490

Doyle, A. C., Smith, E., & Paget, S. (1903). The return of sherlock holmes. Sir Isaac Pitman & Sons Limited.

Ershaid, H., Lizarazu, M., McLaughlin, D., Cooke, M., Simantiraki, O., Koutsogiannaki, M., & Lallier, M. (2023). Contributions of listening effort and intelligibility to cortical tracking of speech in adverse listening conditions. Cortex.

Etard, O., & Reichenbach, T. (2019). Neural speech tracking in the theta and in the delta frequency band differentially encode clarity and comprehension of speech in noise. Journal of Neuroscience, 39(29), 5750–5759.

Fletcher, M. D., Hadeedi, A., Goehring, T., & Mills, S. R. (2019). Electro-haptic enhancement of speech-in-noise performance in cochlear implant users. Scientific Reports, 9(1), 11428.

Fletcher, M. D., Mills, S. R., & Goehring, T. (2018). Vibro-tactile enhancement of speech intelligibility in multi-talker noise for simulated cochlear implant listening. Trends in Hearing, 22, 2331216518797838.

Fu, Q.-J., & Nogaki, G. (2005). Noise susceptibility of cochlear implant users: The role of spectral resolution and smearing. Journal of the Association for Research in Otolaryngology, 6, 19–27.

Fuglsang, S. A., Dau, T., & Hjortkjær, J. (2017). Noise-robust cortical tracking of attended speech in real-world acoustic scenes. NeuroImage, 156, 435–444.

Geirnaert, S., Vandecappelle, S., Alickovic, E., De Cheveigne, A., Lalor, E., Meyer, B. T., … & Bertrand, A. (2021). Electroencephalography-based auditory attention decoding: Toward neurosteered hearing devices. IEEE Signal Processing Magazine, 38(4), 89–102.

Gillis, M., Vanthornhout, J., & Francart, T. (2023). Heard or understood? neural tracking of language features in a comprehensible story, an incomprehensible story and a word list. Eneuro, 10(7).

Gillis, M., Vanthornhout, J., Simon, J. Z., Francart, T., & Brodbeck, C. (2021). Neural markers of speech comprehension: Measuring eeg tracking of linguistic speech representations, controlling the speech acoustics. Journal of Neuroscience, 41(50), 10316–10329.

Gillis, M., Van Canneyt, J., Francart, T., & Vanthornhout, J. (2022). Neural tracking as a diagnostic tool to assess the auditory pathway. Hearing Research, 426, 108607.

Goehring, T., Archer-Boyd, A. W., Arenberg, J. G., & Carlyon, R. P. (2021). The effect of increased channel interaction on speech perception with cochlear implants. Scientific Reports, 11(1), 10383.

Goehring, T., Arenberg, J. G., & Carlyon, R. P. (2020). Using spectral blurring to assess effects of channel interaction on speech-in-noise perception with cochlear implants. Journal of the Association for Research in Otolaryngology, 21, 353– 371.

Goehring, T., Keshavarzi, M., Carlyon, R. P., & Moore, B. C. (2019). Using recurrent neural networks to improve the perception of speech in non-stationary noise by people with cochlear implants. The Journal of the Acoustical Society of America, 146(1), 705–718.

Golumbic, E. M. Z., Ding, N., Bickel, S., Lakatos, P., Schevon, C. A., McKhann, G. M., Goodman, R. R., Emerson, R., Mehta, A. D., Simon, J. Z., et al. (2013). Mechanisms underlying selective neuronal tracking of attended speech at a “cocktail party”. Neuron, 77(5), 980–991.

Grange, J. A., Culling, J. F., Harris, N. S., & Bergfeld, S. (2017). Cochlear implant simulator with independent representation of the full spiral ganglion. The Journal of the Acoustical Society of America, 142(5), EL484–EL489.

Gransier, R., Hofmann, M., van Wieringen, A., & Wouters, J. (2021). Stimulus-evoked phase-locked activity along the human auditory pathway strongly varies across individuals. Scientific Reports, 11(1), 143.

Guérit, F., Deeks, J. M., Arzounian, D., Gransier, R., Wouters, J., & Carlyon, R. P. (2023). Using interleaved stimulation and eeg to measure temporal smoothing and growth of the sustained neural response to cochlear-implant stimulation. Journal of the Association for Research in Otolaryngology, 24(2), 253–264.

Halekoh, U., & Højsgaard, S. (2014). A kenward-roger approximation and parametric bootstrap methods for tests in linear mixed models – the R package pbkrtest. Journal of Statistical Software, 59(9), 1–30. https://www.jstatsoft.org/v59/i09/

Hartig, F. (2017). Package ‘dharma’. https://cran.r-project.org/web/packages/DHARMa/

Hauswald, A., Keitel, A., Chen, Y.-p., Rösch, S., & Weisz, N. (2022). Degradation levels of continuous speech affect neural speech tracking and alpha power differently. European Journal of Neuroscience, 55(11-12), 3288–3302.

Heald, S. L., & Nusbaum, H. C. (2014). Speech perception as an active cognitive process. Frontiers in Systems Neuroscience, 8, 35.

Herrmann, B., & Butler, B. E. (2021). Hearing loss and brain plasticity: The hyperactivity phenomenon. Brain Structure and Function, 226(7), 2019–2039.

Hertrich, I., Dietrich, S., & Ackermann, H. (2013). Tracking the speech signal–time-locked meg signals during perception of ultra-fast and moderately fast speech in blind and in sighted listeners. Brain and Language, 124(1), 9–21.

Hervais-Adelman, A. G., Davis, M. H., Johnsrude, I. S., Taylor, K. J., & Carlyon, R. P. (2011). Generalization of perceptual learning of vocoded speech. Journal of Experimental Psychology: Human Perception and Performance, 37(1), 283.

Heydebrand, G., Hale, S., Potts, L., Gotter, B., & Skinner, M. (2007). Cognitive predictors of improvements in adults’ spoken word recognition six months after cochlear implant activation. Audiology and Neurotology, 12(4), 254–264.

Holden, L. K., Finley, C. C., Firszt, J. B., Holden, T. A., Brenner, C., Potts, L. G., Gotter, B. D., Vanderhoof, S. S., Mispagel, K., Heydebrand, G., et al. (2013). Factors affecting open-set word recognition in adults with cochlear implants. Ear and Hearing, 34(3), 342.

Holdgraf, C. R., Rieger, J. W., Micheli, C., Martin, S., Knight, R. T., & Theunissen, F. E. (2017). Encoding and decoding models in cognitive electrophysiology. Frontiers in Systems Neuroscience, 11, 61.

Holt, L. L., & Lotto, A. J. (2010). Speech perception as categorization. Attention, Perception & Psychophysics, 72(5), 1218–1227.

Howard, M. F., & Poeppel, D. (2010). Discrimination of speech stimuli based on neuronal response phase patterns depends on acoustics but not comprehension. Journal of Neurophysiology, 104(5), 2500–2511.

Intartaglia, B., Zeitnouni, A. G., & Lehmann, A. (2022). Recording eeg in cochlear implant users: Guidelines for experimental design and data analysis for optimizing signal quality and minimizing artifacts. Journal of Neuroscience Methods, 375, 109592.

Iotzov, I., & Parra, L. C. (2019). Eeg can predict speech intelligibility. Journal of Neural Engineering, 16(3), 036008.

Jaeger, B. C., Edwards, L. J., Das, K., & Sen, P. K. (2017). An r 2 statistic for fixed effects in the generalized linear mixed model. Journal of Applied Statistics, 44(6), 1086–1105.

Karunathilake, I. M., Kulasingham, J. P., & Simon, J. Z. (2023). Neural tracking measures of speech intelligibility: Manipulating intelligibility while keeping acoustics unchanged. Proceedings of the National Academy of Sciences, 120(49). 10.1073/pnas.2309166120

Kleiner, M., Brainard, D., Pelli, D., Ingling, A., Murray, R., Broussard, C., & Cornelissen, F. (2007). What is new in psychtoolbox 3. Perception, 36(14), 1–16.

Klug, M., & Kloosterman, N. A. (2022). Zapline-plus: A zapline extension for automatic and adaptive removal of frequency-specific noise artifacts in m/eeg. Human Brain Mapping, 43(9), 2743–2758.

Kong, Y.-Y., Somarowthu, A., & Ding, N. (2015). Effects of spectral degradation on attentional modulation of cortical auditory responses to continuous speech. Journal of the Association for Research in Otolaryngology, 16, 783–796.

Kösem, A., Dai, B., McQueen, J. M., & Hagoort, P. (2023). Neural tracking of speech envelope does not unequivocally reflect intelligibility. NeuroImage, 272, 120040.

Kriegeskorte, N., & Douglas, P. K. (2019). Interpreting encoding and decoding models. Current Opinion in Neurobiology, 55, 167–179.

Lalor, E. C., & Foxe, J. J. (2010). Neural responses to uninterrupted natural speech can be extracted with precise temporal resolution. The European Journal of Neuroscience, 31(1), 189–193. 10.1111/j.1460-9568.2009.07055.x

Lesenfants, D., & Francart, T. (2020). The interplay of top-down focal attention and the cortical tracking of speech. Scientific Reports, 10(1), 6922.

Lesenfants, D., Vanthornhout, J., Verschueren, E., Decruy, L., & Francart, T. (2019). Predicting individual speech intelligibility from the cortical tracking of acoustic- and phonetic-level speech representations. Hearing Research, 380, 1–9.

Lesicko, A. M., & Llano, D. A. (2017). Impact of peripheral hearing loss on top-down auditory processing. Hearing Research, 343, 4–13.

Lizarazu, M., Carreiras, M., Bourguignon, M., Zarraga, A., & Molinaro, N. (2021). Language proficiency entails tuning cortical activity to second language speech. Cerebral Cortex, 31(8), 3820–3831.

Lüdecke, D., Ben-Shachar, M. S., Patil, I., Wiernik, B. M., Bacher, E., Thériault, R., & Makowski, D. (2022). Easystats: Framework for easy statistical modeling, visualization, and reporting [R package]. CRAN. https://easystats.github.io/easystats/

Luke, S. G. (2017). Evaluating significance in linear mixed-effects models in r. Behavior Research Methods, 49, 1494–1502.

MacIntyre, A. D., Cai, C. Q., & Scott, S. K. (2022). Pushing the envelope: Evaluating speech rhythm with different envelope extraction techniques. The Journal of the Acoustical Society of America, 151(3), 2002–2026.

MacIntyre, A. D., & Goehring, T. (2023). Effects of spectral degradation on the cortical tracking of the speech envelope. Proc. INTERSPEECH 2023, 5187–5191.

McGarrigle, R., Knight, S., Rakusen, L., Geller, J., & Mattys, S. (2021). Older adults show a more sustained pattern of effortful listening than young adults. Psychology and Aging, 36(4), 504.

McHaney, J. R., Gnanateja, G. N., Smayda, K. E., Zinszer, B. D., & Chandrasekaran, B. (2021). Cortical tracking of speech in delta band relates to individual differences in speech in noise comprehension in older adults. Ear and Hearing, 42(2), 343– 354.

Millman, R. E., Johnson, S. R., & Prendergast, G. (2015). The role of phase-locking to the temporal envelope of speech in auditory perception and speech intelligibility. Journal of Cognitive Neuroscience, 27(3), 533–545.

Myers, B. R., Lense, M. D., & Gordon, R. L. (2019). Pushing the envelope: Developments in neural entrainment to speech and the biological underpinnings of prosody perception. Brain Sciences, 9(3), 70.

Narain, C., Scott, S. K., Wise, R. J., Rosen, S., Leff, A., Iversen, S., & Matthews, P. (2003). Defining a left-lateralized response specific to intelligible speech using fmri. Cerebral Cortex, 13(12), 1362–1368.

Nogueira, W., & Dolhopiatenko, H. (2022). Predicting speech intelligibility from a selective attention decoding paradigm in cochlear implant users. Journal of Neural Engineering, 19(2), 026037.

Oostenveld, R., Fries, P., Maris, E., & Schoffelen, J.-M. (2011). Fieldtrip: Open source software for advanced analysis of meg, eeg, and invasive electrophysiological data. Computational Intelligence and Neuroscience, 2011, 1–9.

O’Sullivan, J. A., Power, A. J., Mesgarani, N., Rajaram, S., Foxe, J. J., Shinn-Cunningham, B. G., Slaney, M., Shamma, S. A., & Lalor, E. C. (2015). Attentional selection in a cocktail party environment can be decoded from single-trial eeg. Cerebral Cortex, 25(7), 1697–1706.

Oxenham, A. J., & Kreft, H. A. (2014). Speech perception in tones and noise via cochlear implants reveals influence of spectral resolution on temporal processing. Trends in Hearing, 18, 2331216514553783.

Panela, R. A., Copelli, F., & Herrmann, B. (2024). Reliability and generalizability of neural speech tracking in younger and older adults. Neurobiology of Aging, 134, 165–180.

Peelle, J. E., Gross, J., & Davis, M. H. (2013). Phase-locked responses to speech in human auditory cortex are enhanced during comprehension. Cerebral Cortex, 23(6), 1378–1387.

Price, C. N., & Bidelman, G. M. (2021). Attention reinforces human corticofugal system to aid speech perception in noise. NeuroImage, 235, 118014.

Prinsloo, K. D., & Lalor, E. C. (2022). General auditory and speech-specific contributions to cortical envelope tracking revealed using auditory chimeras. Journal of Neuroscience, 42(41), 7782–7798.

R Core Team. (2021). R: A language and environment for statistical computing. R Foundation for Statistical Computing. Vienna, Austria. https://www.R-project.org/

Ramus, F., Dupoux, E., & Mehler, J. (2003). The psychological reality of rhythm classes: Perceptual studies. Proceedings of the 15th International Congress of Phonetic Sciences, 3, 337–342.

Reetzke, R., Gnanateja, G. N., & Chandrasekaran, B. (2021). Neural tracking of the speech envelope is differentially modulated by attention and language experience. Brain and Language, 213, 104891.

Rimmele, J. M., Golumbic, E. Z., Schröger, E., & Poeppel, D. (2015). The effects of selective attention and speech acoustics on neural speech-tracking in a multi-talker scene. Cortex, 68, 144–154.

Samuel, A. G., & Kraljic, T. (2009). Perceptual learning for speech. *Attention, Perception*, & Psychophysics, 71(6), 1207–1218.

Schmidt, F., Chen, Y.-P., Keitel, A., Rösch, S., Hannemann, R., Serman, M., Hauswald, A., & Weisz, N. (2021). Neural speech tracking shifts from the syllabic to the modulation rate of speech as intelligibility decreases. *Psychophysiology*, e14362.

Schmitt, R., Meyer, M., & Giroud, N. (2022). Better speech-in-noise comprehension is associated with enhanced neural speech tracking in older adults with hearing impairment. Cortex, 151, 133–146.

Sokoliuk, R., Degano, G., Melloni, L., Noppeney, U., & Cruse, D. (2021). The influence of auditory attention on rhythmic speech tracking: Implications for studies of unresponsive patients. Frontiers in Human Neuroscience, 15, 702768.

Somers, B., Verschueren, E., & Francart, T. (2018). Neural tracking of the speech envelope in cochlear implant users. Journal of Neural Engineering, 16(1), 016003.

Song, J., & Iverson, P. (2018). Listening effort during speech perception enhances auditory and lexical processing for non-native listeners and accents. Cognition, 179, 163–170.

Svirsky, M. A., Capach, N. H., Neukam, J. D., Azadpour, M., Sagi, E., Hight, A. E., Glassman, E. K., Lavender, A., Seward, K. P., Miller, M. K., et al. (2021). Valid acoustic models of cochlear implants: One size does not fit all. Otology & Neurotology, 42(10), S2–S10.

Terreros, G., & Delano, P. H. (2015). Corticofugal modulation of peripheral auditory responses. Frontiers in Systems Neuroscience, 9, 134.

Thaler, L., Schütz, A. C., Goodale, M. A., & Gegenfurtner, K. R. (2013). What is the best fixation target? the effect of target shape on stability of fixational eye movements. Vision Research, 76, 31–42.

Van Hirtum, T., Somers, B., Dieudonné, B., Verschueren, E., Wouters, J., & Francart, T. (2023). Neural envelope tracking predicts speech intelligibility and hearing aid benefit in children with hearing loss. Hearing Research, 439, 108893.

Vanthornhout, J., Decruy, L., & Francart, T. (2019). Effect of task and attention on neural tracking of speech. Frontiers in Neuroscience, 13, 977.

Vanthornhout, J., Decruy, L., Wouters, J., Simon, J. Z., & Francart, T. (2018). Speech intelligibility predicted from neural entrainment of the speech envelope. Journal of the Association for Research in Otolaryngology, 19, 181–191.

Verschueren, E., Somers, B., & Francart, T. (2019). Neural envelope tracking as a measure of speech understanding in cochlear implant users. Hearing Research, 373, 23–31.

Verschueren, E., Vanthornhout, J., & Francart, T. (2021). The effect of stimulus intensity on neural envelope tracking. Hearing Research, 403, 108175.

Winn, M. B., Edwards, J. R., & Litovsky, R. Y. (2015). The impact of auditory spectral resolution on listening effort revealed by pupil dilation. Ear and Hearing, 36(4), e153.

Xu, N., Zhao, B., Luo, L., Zhang, K., Shao, X., Luan, G., Wang, Q., Hu, W., & Wang, Q. (2023). Two stages of speech envelope tracking in human auditory cortex modulated by speech intelligibility. Cerebral Cortex, 33(5), 2215–2228.

Yasmin, S., Irsik, V. C., Johnsrude, I. S., & Herrmann, B. (2023). The effects of speech masking on neural tracking of acoustic and semantic features of natural speech. Neuropsychologia, 186, 108584.

Zeng, F.-G. (2022). Celebrating the one millionth cochlear implant. JASA Express Letters, 2(7).

Zinszer, B. D., Yuan, Q., Zhang, Z., Chandrasekaran, B., & Guo, T. (2022). Continuous speech tracking in bilinguals reflects adaptation to both language and noise. Brain and Language, 230, 105128.

